# Waves of sumoylation support transcription dynamics during adipocyte differentiation

**DOI:** 10.1101/2021.02.20.432084

**Authors:** Xu Zhao, Ivo A. Hendriks, Stéphanie Le Gras, Tao Ye, Aurélie Nguéa P, Lucia Ramos-Alonso, Guro Flor Lien, Arne Klungland, Bernard Jost, Jorrit M. Enserink, Michael L. Nielsen, Pierre Chymkowitch

**Affiliations:** Department of Biosciences, Faculty of Mathematics and Natural Sciences, University of Oslo, P.O. Box 1066 Blindern, 0316 Oslo, Norway; Department of Microbiology, Oslo University Hospital, Rikshospitalet, 0372 Oslo, Norway; Proteomics Program, Novo Nordisk Foundation Center for Protein Research, Faculty of Health and Medical Sciences, University of Copenhagen, Blegdamsvej 3B, 2200, Copenhagen, Denmark.; Institut de Génétique et de Biologie Moléculaire et Cellulaire, CNRS UMR7104, Inserm U1258, Université de Strasbourg, Illkirch, France; Department of Molecular Cell Biology, Institute for Cancer Research, The Norwegian Radium Hospital, Montebello, 0379 Oslo, Norway; Centre for Cancer Cell Reprogramming, Institute of Clinical Medicine, Faculty of Medicine, University of Oslo, Oslo, Norway

**Keywords:** Adipogenesis, cell differentiation, SUMO, transcription, chromatin

## Abstract

Tight control of gene expression networks required for adipose tissue formation and plasticity is essential for adaptation to energy needs and environmental cues. However, little is known about the mechanisms that orchestrate the dramatic transcriptional changes leading to adipocyte differentiation. We investigated the regulation of nascent transcription by the sumoylation pathway during adipocyte differentiation using SLAMseq and ChIPseq. We discovered that the sumoylation pathway has a dual function in differentiation; it supports the initial downregulation of pre-adipocyte-specific genes, while it promotes the establishment of the mature adipocyte transcriptional program. By characterizing sumoylome dynamics in differentiating adipocytes by mass spectrometry, we found that sumoylation of specific transcription factors like Ppar*γ*/RXR and their co-factors is associated with the transcription of adipogenic genes. Our data demonstrate that the sumoylation pathway coordinates the rewiring of transcriptional networks required for formation of functional adipocytes. This study also provides an in-depth resource of gene transcription dynamics, SUMO-regulated genes and sumoylation sites during adipogenesis.

## Introduction

Posttranslational modification by the small ubiquitin-like modifier SUMO is a conserved, essential and versatile process playing a critical yet poorly understood role in cell differentiation, identity, growth and adaptation to various stimuli (Chymkowitch et al., 2015b; Enserink, 2015; Seeler and Dejean, 2017). Mammals express three major SUMO isoforms: SUMO-1, −2 and −3. SUMO-2 and SUMO-3 are nearly identical and are usually referred to as SUMO-2/3. SUMO precursors are matured by SUMO-specific proteases (SENPs), after which they linked to the E1 activating enzyme SAE1/SAE2. SUMO is then transferred to the E2 conjugating enzyme Ubc9, which conjugates SUMO to lysine residues of target proteins in a process that is often facilitated by E3 ligases. Sumoylation can alter the stability, conformation, interactions, or subcellular localization of target proteins. SUMO targets can be desumoylated by SENPs (Chymkowitch et al., 2015b).

Proteomic studies have shown that most SUMO targets are involved in chromatin regulation, chromosome integrity, mRNA processing and transcription (Hendriks et al., 2018; Hendriks and Vertegaal, 2016; Paakinaho et al., 2021). Consistently, genome-wide chromatin immunoprecipitation-sequencing (ChIPseq) experiments revealed that SUMO is present at intragenic regions, enhancers and transcription factor binding sites (Chymkowitch et al., 2015a; Chymkowitch et al., 2017; Cossec et al., 2018; Neyret-Kahn et al., 2013). The SUMO chromatin landscape is substantially remodeled in response to heat shock, inflammation, oncogene-induced senescence, nutrient deprivation and pro-growth signals (Chymkowitch et al., 2015a; Chymkowitch et al., 2017; Decque et al., 2016; Neyret-Kahn et al., 2013; Nguea et al., 2019; Niskanen et al., 2015).

Recent studies have revealed that SUMO is also important for cellular identity (Cossec et al., 2018; Theurillat et al., 2020). However, these studies only compared undifferentiated versus terminally differentiated cells, precluding the sumoylation dynamics that may occur throughout the differentiation process. Therefore, there exists a need for a biologically relevant model to study sumoylome dynamics during differentiation.

Interestingly, mouse knockout studies have revealed that animals lacking Sumo-1 or Senp2 are resistant to high fat diet (HFD)-induced obesity (Mikkonen et al., 2013; Zheng et al., 2018), whereas specific loss of *UBC9* in white adipose tissue triggers lipoatrophy (Cox et al., 2020). Consistently, loss of Sumo-1, Ubc9, Senp1 or Senp2 in cellular models impedes adipogenesis and is associated with altered expression of genes under the control of PPAR*γ,* cEBP*α*, or cEBP*δ,* as well as genes involved in lipid and energy metabolism (Chung et al., 2010; Cignarelli et al., 2010; Liu et al., 2014). This suggests that formation of adipose tissue depends on the sumoylation pathway, although the dynamics of this process remain very poorly described.

In this study, we used adipocyte differentiation (AD) as a model system to uncover the dynamics of the sumoylome, the SUMO-chromatin landscape and SUMO-dependent gene transcription during differentiation. Our data show that SUMO plays a dual role during adipogenesis by supporting the transcription of pre-adipocyte genes and promoting adipogenic genes transcription to establish adipocyte identity.

## Results

### Inhibition of sumoylation triggers lipoatrophy

To begin investigating the role of the sumoylation pathway during adipogenesis, we made use of the mouse 3T3-L1 cell model (Green and Kehinde, 1975). Incubation of these cells in differentiation medium induces a well-defined adipocyte differentiation process, during which cells transition from a pre-adipocyte (PA) stage via a clonal expansion (CE) stage into mature adipocytes (MA). We induced differentiation of 3T3-L1 cells either in absence or in presence of the selective SAE1/2 inhibitor ML-792, which does not inhibit other ubiquitin family pathways (He et al., 2017)(Fig. S1A). Western blotting of control cells with a SUMO2/3 antibody revealed a progressive increase in SUMO conjugates during AD (Fig. S1B), indicating mobilization of the sumoylation pathway, whereas treatment with ML-792 almost completely prevented the accumulation of SUMO conjugates (Fig. S1B).

Next, we assessed the effect of ML-792 on fat accumulation using Bodipy reagents. Seven days after adipogenic induction there was a normal accumulation of lipid droplets in control cells (Fig. S1C), which is a key feature of adipocyte differentiation (Green and Kehinde, 1975). In contrast, treatment with ML-792 resulted in lipoatrophy (Fig. S1C), which was characterized by numerous small lipid droplets (Fig. S1D-E), mirroring the recently reported phenotype of loss of Ubc9 in mouse white adipose tissue (Cox et al., 2020). These data show that an active sumoylation pathway is required for efficient adipogenesis.

### Mapping the nascent transcriptome landscape of differentiating adipocytes

Given that sumoylation mostly targets nuclear proteins involved in chromatin and gene regulation (Hendriks and Vertegaal, 2016), we hypothesized that ML-792 treatment resulted in lipoatrophy due to defects in adipogenic gene expression programs. We decided to use SLAMseq (Herzog et al., 2017) to specifically analyze the transcriptome-wide kinetics of nascent RNA synthesis and turnover during AD (Fig. 1A). This revealed strong transcriptional differences between day −2, day 1, and day 3, whereas day 7 showed transcriptional levels similar to day 3 (Fig. 1B and Supp. Fig. S1F). This indicates that most of the transcriptional rewiring during AD occurs until day 3, after which transcriptional output stabilizes as the adipocyte matures.

**Fig. 1.**
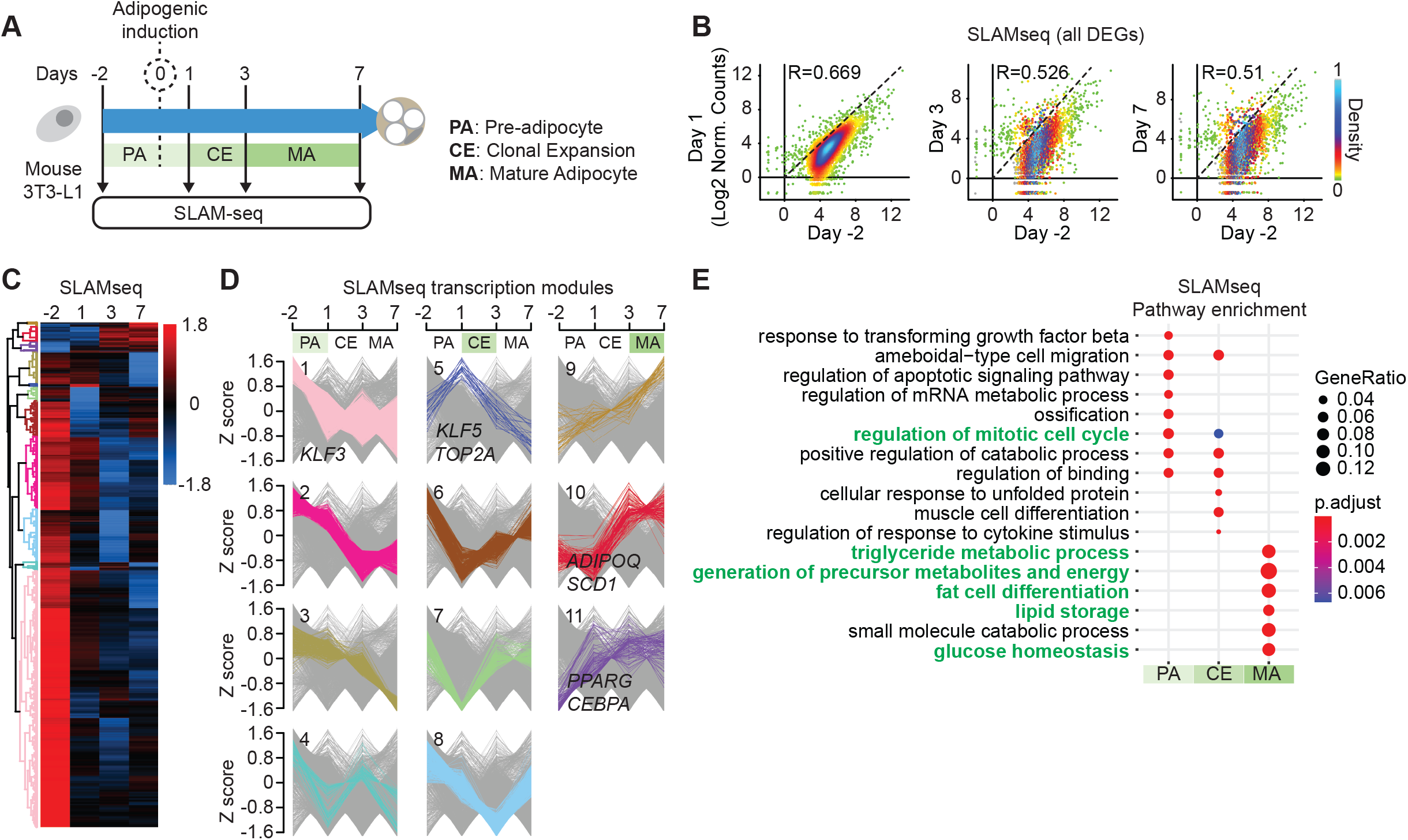
Characterization of nascent transcriptome dynamics during adipocyte differentiation. (A) Experimental layout of time-course experiments. (B) Scatter plot showing the nascent transcription levels for all time-course differentially expressed (DE) transcripts at each time point vs Day-2. The nascent transcription levels are presented as log2 transformed normalized counts. R, Pearson correlation coefficient. (C and D) Hierarchical clustering of DE transcripts (*C*). Clusters are categorized to 3 stage-specific modules (*E*). PA: pre-adipocyte, CE: clonal expansion, MA: mature adipocyte. Z score is calculated for log2 transformed normalized counts. (E) GO analysis of DEGs in stage-specific modules.

In total we identified 3705 DEGs during AD (Table S1 and STAR methods). Our data clearly show a dramatic general downregulation of transcription upon adipogenic induction (Fig. 1B). Clustering analysis revealed 11 transcriptional modules (Fig. 1C and D). The vast majority of genes (3433 out of 3705, comprising modules 1-4 and 6-8) were transcribed in the PA stage and then downregulated upon adipogenic induction. Transcription of these genes reached minimum levels either during CE or at MA stage (Fig. 1C and D, Fig. S1G and Table S2), such as the anti-adipogenic gene *KLF3* (Garcia-Nino and Zazueta, 2021). Interestingly, we noticed that a small number of 273 genes displayed increased transcription upon adipogenic induction. Transcription of a subset of these genes (module 5) was highly dynamic, showing a sharp increase during the transition from PA to CE, followed by strong downregulation as cells enter the MA stage. Closer inspection revealed that this module contained genes involved in mitotic cell cycle regulation and adipogenesis, such as *TOP2A* and *KLF5* (Garcia-Nino and Zazueta, 2021) (Fig. 1C and D, Fig. S1G and Table S2). Other genes showed a more gradual increase in transcription levels and their expression remained high during later stages of AD (modules 9, 10 and 11), such as *PPARG* and *ADIPOQ*, both known to be highly expressed in MAs (Fig. 1C and D, Fig. S1G and Table S2).

Pathway enrichment analysis of SLAMseq DEGs revealed a partial overlap between terms in PA and CE (Fig. 1E). These common terms notably included mitotic regulation of the cell cycle, indicating that transcriptional upregulation of cell cycle genes required for CE is one of the earliest event following adipogenic induction. In contrast, MA-specific transcription modules 9-11 featured a strong enrichment in genes involved in fat cell differentiation and mature adipocyte functions, including triglyceride metabolism, glucose homeostasis, and lipid storage (Fig. 1E).

### SUMO promotes transcription of pro-adipogenic genes

Next, we studied the effect of the sumoylation pathway on the transcriptional landscape of differentiating 3T3-L1 cells by treating cells with ML-792 (Fig. 2A). Differential expression analysis of ML-792 versus DMSO control samples revealed 905 DEGs during AD (1334 transcripts; Table S6, Table S3 and STAR methods). Among them, 636 DEGs (927 DE transcripts) were significantly regulated over the normal time course (Fig. S2A). Pathway enrichment analysis of the common DEGs revealed various processes including regulation of mitosis and fat cell differentiation, the latest being enriched beyond normal expectations (Fig. S2B and C). We found that 407 genes were uniquely affected by ML792 treatment, and hierarchical clustering revealed that sumoylation regulates the transcription of many genes located in PA, CE and MA modules (Fig. 2B and C and Table S4). Pearson correlation showed that PA modules were only mildly affected by ML-792 (Fig. 2D and S2D), but that CE DEGs and in particular MA DEGs showed a strong deviation from transcription patterns observed in the control experiment (Fig. 2D and S2D). This indicates that sumoylation is important for establishing the mature adipocyte program.

**Fig. 2.**
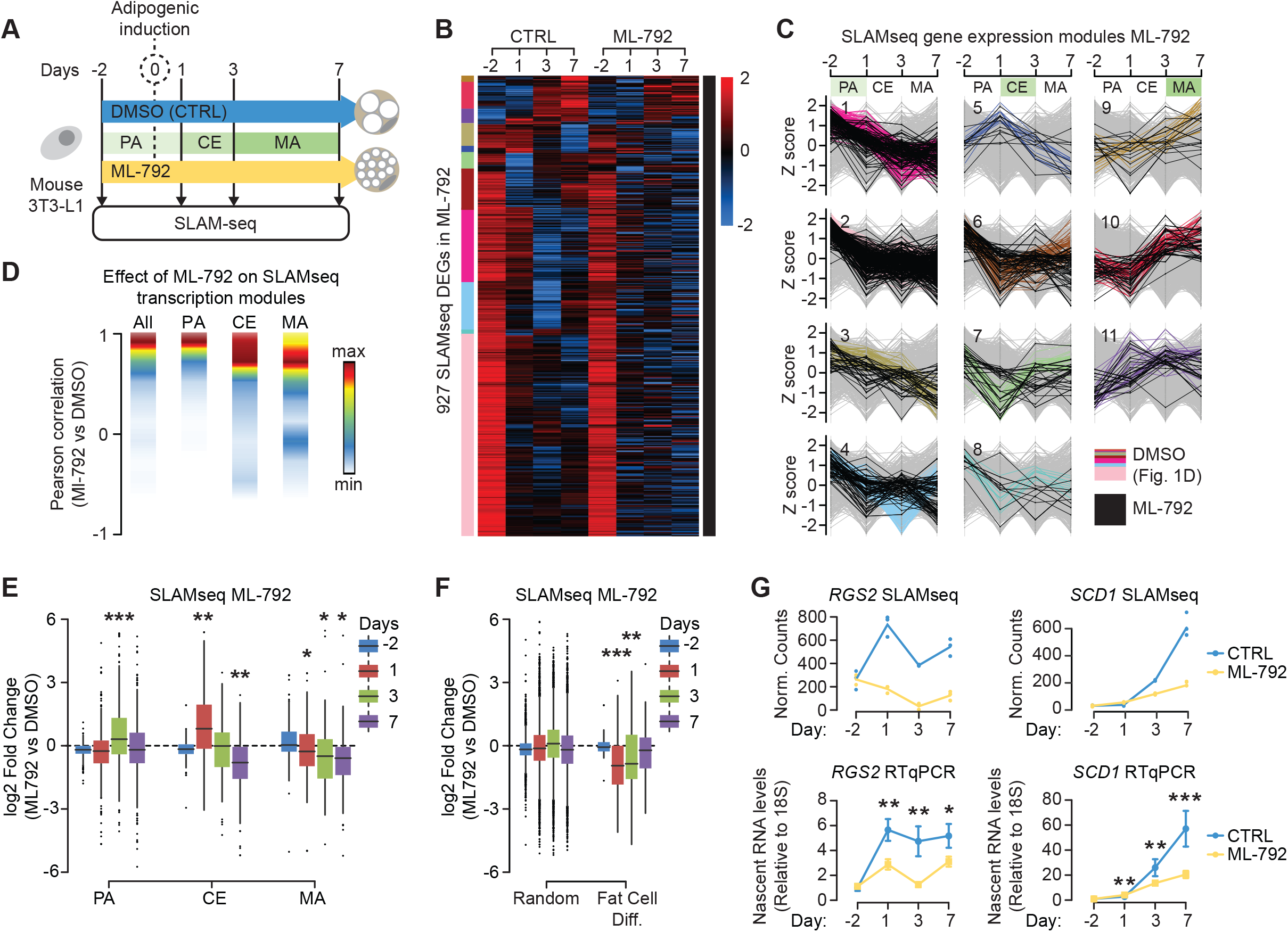
Sumoylation supports adipogenic genes transcription. (A) Experimental setup of time-course SLAMseq experiments with and without ML-792 treatment. (B) Heatmap showing ML792-treated DE transcripts in the time-course clusters *(see* Fig. 1D). CTRL: DMSO. (C) Clustering profiles showing the nascent transcription levels for ML792-treated DE transcripts with and without ML-792 treatment in stage-specific modules. Z scores are calculated for log2 transformed normalized counts. (D) Density plot showing the distributions of Pearson correlation coefficients between the transcription profiles of DE transcripts in DMSO and ML-792 conditions in stage-specific modules. The y axis, presented as Pearson correlation coefficient, is segmented into 10 bins. The number of transcripts within each bin is presented as a color code. (E and F) Boxplots showing effects of ML-792 on the transcription of DE transcripts in stage-specific modules *(E)*, fat cell differentiation term and randomly picked population *(F).* The significance of mean comparison is determined by paired t-test. *: p ≤ 0.05, **: p ≤ 0.01, ***: p ≤ 0.001. (G) Loess regression lines showing transcription profiles of representative DE transcripts in DMSO and ML-792 conditions. The y axis is presented as normalized counts in SLAMseq. DMSO, red; ML-792, green. (H) Validation of transcription profiles of representative DE transcripts in DMSO and ML-792 conditions by RT-qPCR. Data are presented as mean ± standard deviation (SD) (ML-792 vs DMSO). DMSO, blue; ML-792, yellow.

We next dissected the effect of sumoylation on AD modules by assessing transcriptional deregulation in each module for each time point. Interestingly, at day 3 many PA genes had failed to be downregulated when cells were treated with ML792, indicating that the sumoylation pathway is important for repressing PA genes upon adipogenic induction (Fig. 2E). We also found that ML792 treatment caused many CE genes to remain strongly upregulated at day 1 but to become downregulated at day 7 (Fig. 2E). This suggests that SUMO has a dual role in controlling these genes; it first represses these genes during entry into CE and then it reactivates them when cells enter MA stage. Finally, there was a substantial group of MA genes that failed to become activated in presence of ML-792 (Fig. 2E), showing that sumoylation supports transcription in mature adipocytes. More specifically, ML-792 treatment resulted in downregulation of MA-specific genes involved in fat cell differentiation starting from day 1 after adipogenic induction, while transcription of an equal number of randomly selected control genes was not affected (Fig. 2F and Table S5). The inhibitory effect of ML-792 on the transcription of adipogenic genes was validated for *SCD1* and *RGS2* by qPCR (Fig. 2G) (Bai et al., 2015; Nishizuka et al., 2001).

Taken together, these data indicate that sumoylation has a complex and dynamic function in regulating the adipocyte differentiation program. In the early stages of AD it is important for repression of PA genes and activation of CE genes, whereas in later stages it is required for maintaining high transcription levels of genes involved in fat cell differentiation and adipogenic function in mature adipocytes.

### Chromatin-bound SUMO supports the transcriptional identity switch from pre-adipocyte to mature adipocyte

Previous studies showed that the SUMO-chromatin landscape is highly dynamic in response to various signals (Chymkowitch et al., 2015a; Chymkowitch et al., 2017; Cossec et al., 2018; Niskanen et al., 2015; Paakinaho et al., 2021). To gain insight into the dynamics and location of SUMO at chromatin during AD we performed ChIPseq experiments using an anti Sumo-2/3 antibody (Fig. 3A). PCA revealed good correlation between biological replicates and strong variation between time points (Fig. S3A). We identified 35,659 SUMO peaks (Table S6), which were primarily located at transcription units, especially at promoters (Fig. S3B). Strikingly, both the number and intensity of SUMO peaks increased after induction of adipogenesis (Fig. 3B). 1850 significant differential SUMO binding sites were identified over the time course (Table S7). Pathway enrichment analysis showed that SUMO peaks are overrepresented at genes involved in fat cell differentiation and genes with hallmark MA functions, such as lipid metabolism (Fig. S3C).

**Fig. 3.**
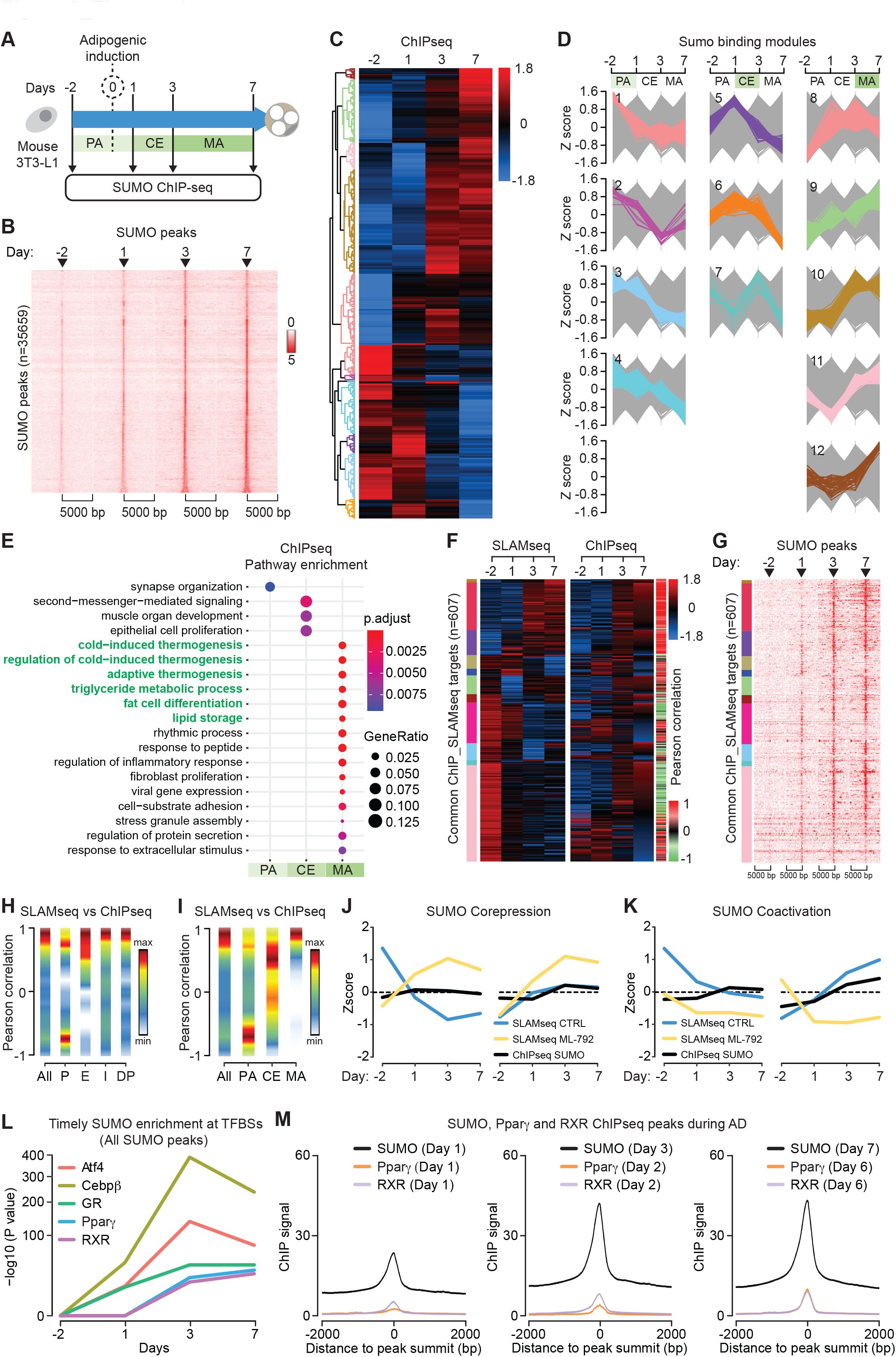
A dual role for SUMO at the chromatin. (A) Experimental setup of time-course ChIPseq experiment. (B) Heatmaps of SUMO occupancy within a 10 kb window centered on SUMO peak summit. SUMO peaks are sorted by increasing size. The intensity represents log10 transformed FPKM. (C and D) Hierarchical clustering of differentially enriched SUMO peaks during the time course(*C*). Clusters are further categorized to 3 stage-specific modules: PA, CE and MA (*D*). Z scores are calculated for log2 transformed normalized counts. (E) GO analysis of SUMO target genes in PA, CE and MA modules. (F) Heatmaps of the time course clusters (see Fig. 1D) and corresponding SUMO peaks. Pearson correlation coefficients between transcription levels and SUMO occupancy of genes are indicated by gradient color. (G) Heatmaps of SUMO occupancy within 10 kb window centered on SUMO summits. SUMO peaks are sorted in the same order as *(F)*. The intensity represents log10 transformed FPKM. (H and I) Distributions of Pearson correlation coefficients in *(F)* in groups categorized base on genomic locations of SUMO peaks *(H)* and in three AD modules *(I)*. All: all SUMO peaks, P: promoter-TSS, E: exon, I: intron, DP: distant promoter. The y axis, presented as Pearson correlation coefficient, is segmented into 10 bins. The number of transcripts within each bin is presented as a color code. (J and K) Profiles (Loess regression lines) of SLAMseq (CTRL, blue; ML-792, yellow) and SUMO ChIPseq (Black) for up-regulated genes *(J, correppression)* and down-regulated genes *(K, coactivation)* upon ML-792 treatment. Z scores are calculated for log2 transformed normalized counts. (L) Motif enrichment of SUMO binding at TFBSs during AD. P value accounts for motif enrichment. (M) Plots of SUMO, Pparγ and RXR ChIPseq signals within 4 kb window centered on peak summits during AD.

Hierarchical clustering identified SUMO binding modules with highest ChIP signals at PA, CE and MA stages (Fig. 3C-D and Table S7). PA modules 1-4 contained 544 SUMO peaks of which the intensity decreased upon adipogenic induction. These peaks were assigned to a heterogeneous set of genes that did not show significant pathway enrichment (Fig. 3E). The same was observed for CE modules 5-7 that contained 170 peaks with highest intensity at day 1 or 3 (Fig. 3E). Finally, MA modules 8-12 contained 1136 peaks with highest intensity in MAs and a very strong enrichment for genes involved in adipogenic functions as well as adaptive thermogenesis (Fig. 3E). These data indicate that chromatin-bound SUMO may support adipocyte function and metabolism via transcriptional regulation.

By comparing the SLAMseq transcription modules identified in Fig. 1D with SUMO ChIPseq data, followed by hierarchical clustering, we found that distinct peaks of SUMO were present both at repressed genes and at activated genes upon induction of adipogenic differentiation (Fig. 3F and G and Table S8). Pearson scoring revealed a general positive correlation between transcription and the presence of SUMO (Fig. 3H-I), which was especially true for exons, introns and distant promoters (Fig. 3H). However, the presence of SUMO at promoter regions both positively and negatively correlated with transcription (Fig. 3H). Most PA genes, including the AD inhibitor *KLF10*, appeared to be repressed in presence of SUMO upon induction of adipogenic differentiation (Fig. 3I and Fig. S3D). In clear contrast, the presence of SUMO at CE genes and in particular at MA genes, such as the adipogenic gene *FABP4*, showed a positive correlation with transcription (Fig. 3I and Fig. S3D).

Based on these findings, we hypothesized that adipogenic stimulation on the one hand leads to increased sumoylation of transcription factors to inhibit transcription of PA genes, while on the other hand it increases transcription of MA genes. We integrated ML-792 SLAMseq data with CTRL SLAMseq and ChIPseq datasets by hierarchical clustering (Fig. S3E and Table S9). This revealed two main categories of SUMO-target genes independently of whether they are activated or repressed upon adipogenic induction: Genes at which SUMO functions as a corepressor, and genes where it serves as a coactivator (Fig. 3J and K and Table S10). Generally, genes activated by SUMO are involved in fat cell differentiation and carbohydrate metabolism, whereas genes repressed by SUMO are not related to adipogenesis (Fig. S3F).

We conclude that (i) chromatin-bound SUMO has a dual role during AD by repressing PA genes while promoting MA genes; and (ii) that SUMO plays an instrumental role in the transcriptional identity switch from pre-adipocyte to mature and functional adipocyte.

### Adipogenic differentiation is associated with waves of SUMO on chromatin

To identify potential transcription factor binding sites (TFBS) enriched for SUMO, we performed a motif search using the SUMO ChIPseq dataset as input (Table S11). While no specific TFBS sequence for adipogenic transcription factors was retrieved at day −2 (Fig. 3L and S3G), we observed significant recruitment of SUMO at critical adipogenic TFBSs over time after stimulation of adipogenic differentiation (Fig. 3L and Fig. S3G). A first wave of SUMO was detected at binding sites for Cebp*β*, GR and Atf4 shortly after adipogenic induction from day 1 until day 7. A second wave of SUMO occurred at Ppar*γ* and RXR response elements, starting on day 3 and lasting until day 7. Similar results were obtained with TFBS analysis of SUMO peaks that are found at AD-regulated genes identified in the SLAMseq dataset (Fig. S3H). To validate these findings, we compared our SUMO ChIPseq data with published Ppar*γ*/RXR ChIPseq (Nielsen et al., 2008), and found that there was a very significant and progressive timely overlap between SUMO, Ppar*γ* and RXR peaks during AD (Fig 3L and M and Table S12). Notably, SUMO recruitment to Ppar*γ*/RXR TFBSs followed the known timeline of recruitment of these TFs during AD (Lefterova et al., 2014; Nielsen et al., 2008).

These data show that the increase in SUMO binding to genes that are upregulated during AD occurs at very specific adipogenic TFBS like Cebp*β*, GR and Ppar*γ*/RXR.

### Site-specific characterization of the SUMOylome during adipocyte differentiation

Our ChIPseq data show that SUMO binds to specific TFBSs during AD. To identify adipogenesis-specific SUMO2 substrates, we carried out site-specific characterization of the endogenous SUMOylome by mass spectrometry during AD (Fig. 4A)(Hendriks et al., 2018). PCA of SUMO-modified lysine residues of four independent experiments demonstrated high reproducibility between replicates and considerable differences between the time points (Fig. S4A-B). Across all time points, fractions and biological replicates, we identified 5230 SUMO-modified peptides that mapped to 3706 unique SUMO sites. Out of all SUMO sites, 3137 (∼85%) could be quantified in quadruplicate (Table S13).

**Fig. 4.**
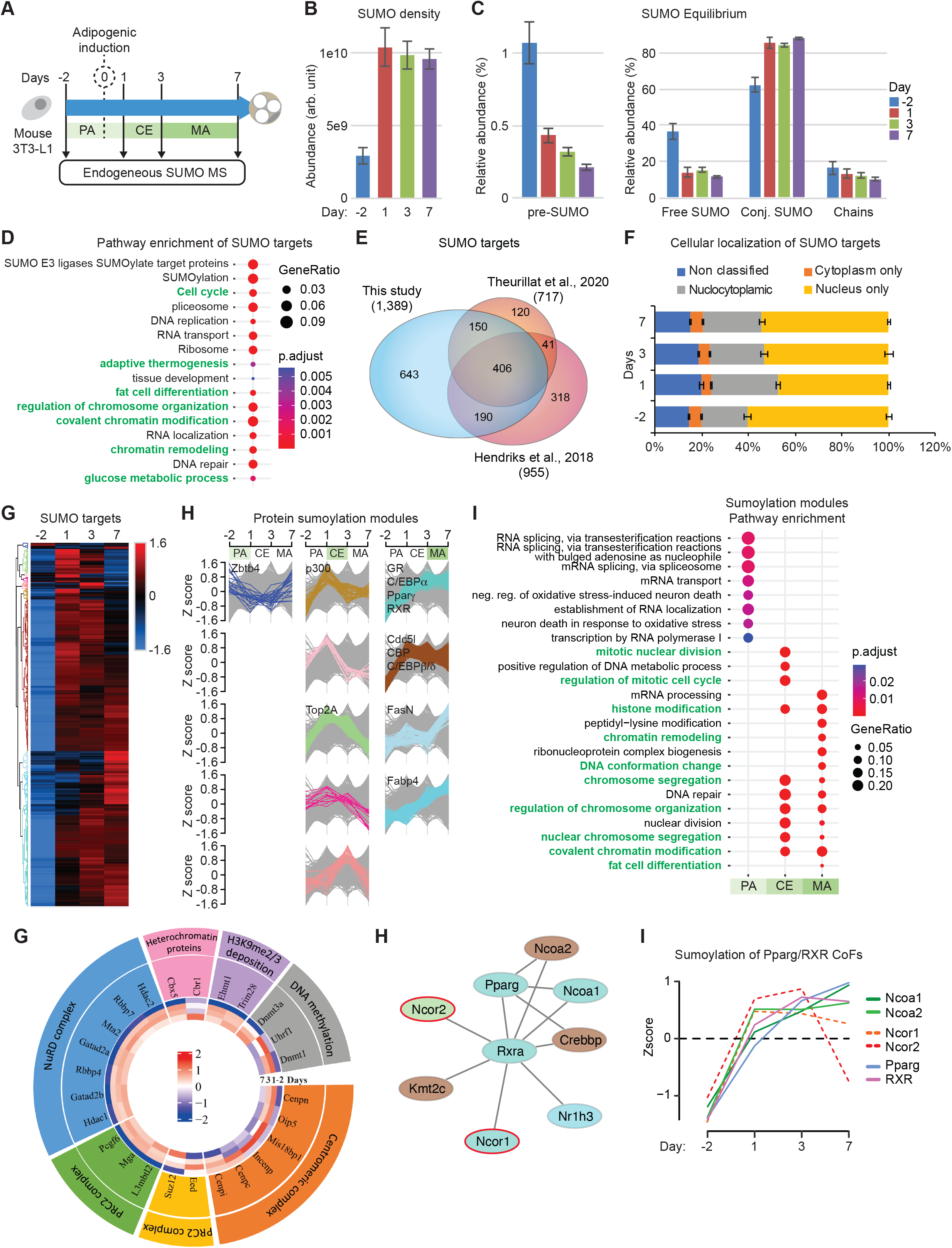
Site specific characterization of the endogenous SUMOylome of differentiating adipocytes. (A) Experimental setup of time-course endogenous SUMO2/3 proteomics. (B) Quantification of total SUMO2/3 density normalized to input amount. (C) The equilibrium for SUMO2/3 in multiple forms. The y axis is presented as relative abundance to total SUMO2/3 density at each time point. (D) GO analysis of SUMO targets. (E) Venn diagram of SUMO targets identified in this study and two recently published studies that used endogeneous detection of sumoylated sites. (F) The percentages of SUMO targets in nucleus and cytoplasm fractions during AD. (G and H) Hierarchical clustering of SUMO targets (*G*). Clusters are categorized in 3 modules: PA, CE and MA (*H*). (I) GO analysis of proteins in sumoylation modules.

In accordance with SUMO western blots (Fig. S1B), MS experiments showed that the overall density of SUMO increases upon adipogenic induction (Fig. 4B). Furthermore, when we analyzed the SUMO equilibrium, i.e. the distribution of SUMO across the entire system and whether it exists in a free or in a conjugated form, we found a larger portion of immature as well as free SUMO prior to induction of differentiation (Fig. 4C). Upon adipogenic induction the consumption of free SUMO increased to reach near-maximum levels at the later time points (Fig. 4C). We did not detect significant alteration of the formation of SUMO chains during AD (Fig. 4C).

In total, we identified 1389 SUMO target proteins of which 1250 could be quantified in quadruplicate with high reproducibility (Table S14, Suppl. Fig. S4C). Pathway enrichment analysis showed that these targets are involved in chromatin regulation, transcription, chromosome maintenance, and cell cycle progression (Fig. 4D and Table S15), with specific enrichment of pathways that are critical for adipose tissue development, function, adaptive thermogenesis, and brown cell differentiation (Fig. 4D and E, Fig S4D and Table S16). Remarkably the majority of SUMO targets (∼80%) are nuclear or nucleocytoplasmic proteins while only ∼5% are cytoplasmic (Fig. 4F and Table S17). This is consistent with previous studies (Hendriks and Vertegaal, 2016) and the sharp and sustained increase of SUMO binding to the chromatin observed in Fig. 3.

Hierarchical clustering of the SUMO targets underlined the sharp remodeling of the SUMOylome occurring after adipogenic induction (Fig. 4G and Table S18). Only very few proteins (21) were highly sumoylated at PA stage; a notable example being the transcriptional repressor Zbtb4 (Fig. 4H and Fig. S4E). 174 out of 1250 SUMO targets became highly sumoylated during CE (Fig. 4H). These proteins are involved in mitotic chromosome and centromere regulation, nucleosome modification and DNA methylation, and include Top2A, CdcA5, p300, Cenps, Dnmt1, Hdac2 and Hdac4 (Fig. 4I-H and Fig. S4E), supporting the idea that SUMO is important for mitotic events during CE. The 1055 MA-specific targets were enriched for proteins critical for transcriptional silencing like polycomb and Dnmt1, chromatin remodeling like NurD, histone modifications like CBP and Trim28 and fat cell differentiation like the pro-adipogenesis TFs GR, cEBP*α/δ*, Ppar*γ* and RXR (Fig. 4-G). These targets also include a few proteins involved in fat metabolism, such as Fabp4 or FasN (Fig. 4I, Fig. S4E). Thus, sumoylation of adipogenic TFs clearly correlates with MA stage and indicates that SUMO supports adipogenic function at the transcription level.

Next, we mapped the interaction network of sumoylated TFs using STRING network analysis (Szklarczyk et al., 2011). Integration of the STRING network with sumoylation modules revealed that two TFs/CoFs were sumoylated in PA, 12 in CE and 96 in MA modules (Fig. S4F). Interestingly, only nine proteins were classified as transcriptional repressors (Fig. S4F). A major interaction node was centered on Cdc5l and interactors, which are sumoylated early during AD and in MAs (Fig. 4H, Fig. S4F and Table S19). Interestingly, Cdc5l is involved in mitotic progression and has been associated with fast cell cycle progression during the early phase of stem cell reprogramming (Jeong et al., 2017), suggesting that sumoylation of Cdc5l and its interactors may contribute to cell cycle progression during CE. Another major node is centered on the histone acetyltransferase Crebbp (CBP), which is an interaction partner and transcriptional co-activator for many TFs, including the nuclear receptors Ppar*γ*/RXR (Fig. 4H and Fig. S4F). Ppar*γ*/RXR response elements are strongly occupied by SUMO in MA stage (Fig. 3L and M), consistent with a model in which sumoylation of Ppar*γ*/RXR and coactivators is important for transcriptional regulation in mature adipocytes (Figure 4I).

These data indicate that the timely sumoylation of mitotic, chromatin and transcriptional regulators supports the CE and MA stages of AD. Furthermore, our data strongly suggest that sumoylation of chromatin bound Ppar*γ*/RXR and their CoFs supports adipogenic transcription.

### Sumoylation supports transcription of Pparγ/RXR target genes in mature adipocytes

Our data indicate that the SUMO peaks detected at Ppar*γ*/RXR binding sites (Fig. 3 and S3) are due to the presence of sumoylated TFs and co-activators, including Ppar*γ*/RXR itself (Fig. 4). We integrated the SUMO MS, SUMO SLAMseq, SUMO ChIPseq, and Ppar*γ*/RXR ChIPseq datasets and focused on Ppar*γ*/RXR target genes of which the regulation is critical during AD (Nielsen et al., 2008). We identified 143 DEGs, both normal time-course DEGs and ML-792 treated DEGs, at which SUMO perfectly overlapped with Ppar*γ*/RXR (Figure 5A and Table S20). Note that these genes are adipogenic genes (Fig. S5A) that are either repressed (PA genes) or activated (MA genes) by adipogenic stimulation (Fig. 5B and S5B). Ppar*γ*/RXR became sumoylated upon adipogenic induction, which was subsequently followed by an increased presence of SUMO at Ppar*γ*/RXR response elements (Fig. S5B *and example of SCD1 gene in Fig S5C*). Importantly, treatment with ML-792 led to the downregulation of PA genes and upregulation of MA genes at day −2, whereas MA genes were strongly downregulated at day 7 (Fig. 5B and C and Fig. S5B and C). These data indicate that the timely regulation of Ppar*γ*/RXR target genes in response to adipogenic stimulation requires the activity of the sumoylation pathway. Inhibiting the sumoylation process perturbed the transcription of these genes and did not allow for development of proper adipogenic functions, which resulted in lipoatrophy (Fig. S1B and 5C).

**Fig. 5.**
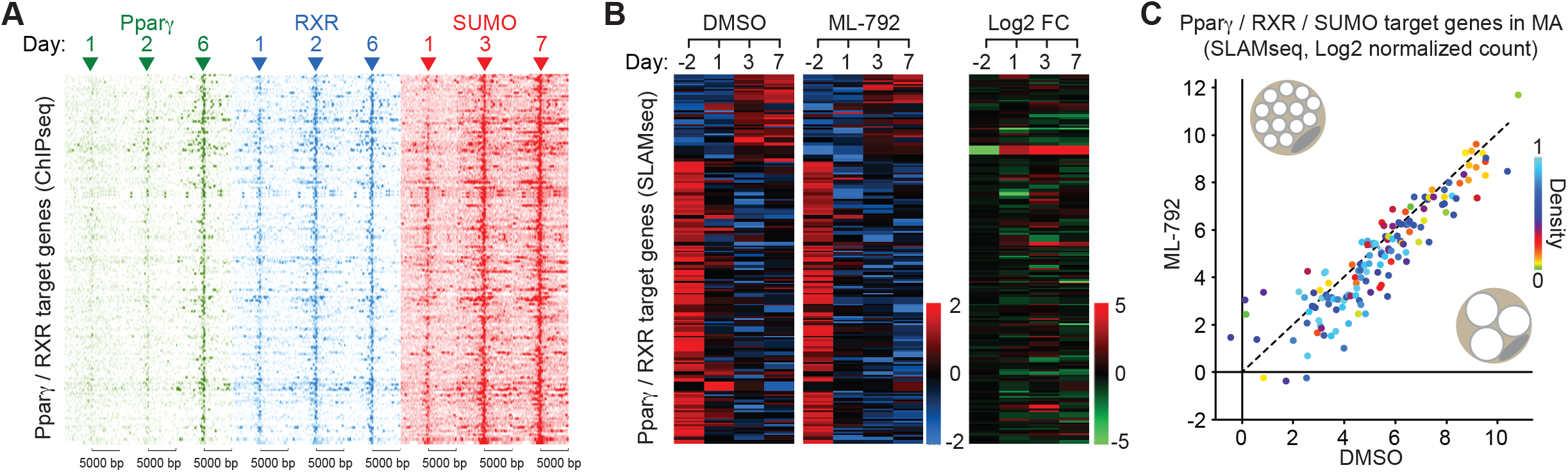
Pparγ, RXR and SUMO promote adipogenic genes transcription. (A) Heatmaps of Pparγ, RXR and SUMO occupancy within a 10 kb window centered on common peak summit of 208 Pparγ/RXR/SUMO common target genes. The intensity represents log10 transformed FPKM. SUMO peaks are sorted as the order of rows in *(B)*. (B) Hierarchical clustering of 208 Pparγ/RXR/SUMO common target genes in *(A)*. Z score is calculated for log2 transformed normalized counts. (C) Scatter plot showing the global downregulation of transcription for 208 Pparγ/RXR/SUMO common target genes upon ML-792 treatment in MA. The nascent transcription levels are presented as log2 transformed normalized counts.

## Discussion

Evidence that the sumoylation pathway supports adipose tissue formation and function has been provided by previous studies (Chung et al., 2010; Cignarelli et al., 2010; Cox et al., 2020; Liu et al., 2014; Mikkonen et al., 2013; Zheng et al., 2018). The majority of these studies have concluded that sumoylation generally inhibits adipogenic transcription factors. This view has been standing for many years, but recent systematic large-scale studies have revealed that SUMO not only represses but can also promote transcription, depending on the cellular system and the physiological context (Boulanger et al., 2021; Chymkowitch et al., 2015b). Systematic analysis of the dynamics of SUMO-regulated gene expression in differentiating adipocytes has never been performed. In the current study we have shown that adipogenesis involves dynamic reorganization of the sumoylome at transcription units, and that SUMO is important for establishment and robustness of the adipogenic transcription program to enforce the identity shift from pre-adipocyte to mature adipocyte.

The presence of SUMO on chromatin is generally believed to be associated with the establishment of repressive chromatin (Cossec et al., 2018; Decque et al., 2016; Neyret-Kahn et al., 2013; Niskanen et al., 2015; Theurillat et al., 2020). Indeed, we found that SUMO has a predominantly repressive function at the early stages of AD, when 92% of PA genes are repressed; treatment with ML-792 results in unscheduled activation of these genes. The mechanism by which SUMO represses these genes is the focus of our ongoing studies. Sumoylation of transcriptional activators could inhibit their transactivation activity via the recruitment of corepressors that close the chromatin, as has recently been shown for the glucocorticoid receptor in HEK-293 cells (Paakinaho et al., 2021). Alternatively, our MS data indicate that SUMO may result in the recruitment or activation of chromatin silencers to establish and maintain PA gene silencing in mature adipocytes, such as the heterochromatin polycomb complex or DNMTs (Cossec et al., 2018; Theurillat et al., 2020).

However, the sumoylation pathway also has a distinct positive effect on transcription of a subset of genes. Here, adipogenic induction triggers the recruitment of SUMO to genes known to be important for the MA state, and the presence of SUMO at these genes correlated positively with their transcription. We observed an unexpected dynamic involvement of SUMO in regulation of transcription during AD, which appears to involve two waves of sumoylation at the promoter of MA genes. The early wave affects Atf4, cEBP*β* and GR binding sites. The second wave of sumoylation mainly affects Ppar*γ*/RXR response elements (PPRE) and peaks at the MA stage. It is possible that multiple components of the Ppar*γ*/RXR complex become sumoylated, because our MS data show that sumoylation of Ppar*γ*/RXR coactivators Ncoa1, Ncoa2 and CBP and chromatin remodeling complexes like NurD are all SUMO targets. Exactly how SUMO regulates these proteins to activate transcription remains to be established, but could involve stabilization of protein-protein interactions within the complex.

Of note, our findings that sumoylation of Ppar*γ*/RXR and its coactivators supports the adipogenic transcription program is in apparent contrast with previous reports showing that sumoylation inhibits Ppar*γ* activity (Floyd and Stephens, 2004; Ohshima et al., 2004; Pascual et al., 2005). However, these previous studies mostly addressed the regulation of Ppar*γ* by SUMO1 rather than SUMO2/3 using single Ppar*γ* mutants and a small selection of model genes (Brunmeir and Xu, 2018). In contrast, our comprehensive genome-wide approach clearly shows that inhibition of the sumoylation pathway strongly reduces the expression of Pparg/RXR target genes. Thus, the physiological relevance of the sumoylation pathway during AD is that it has a net overall positive effect on the Pparg/RXR transcription program, which is also consistent with the lipoatrophy phenotype that we and others have observed upon inhibition of sumoylation(Cox et al., 2020; Mikkonen et al., 2013). A similar apparent discrepancy has been observed in yeast, where one study reported that sumoylation of specific lysines in the RNA polymerase III complex activates the enzyme (Chymkowitch et al., 2017), whereas a second study found that sumoylation of another set of lysines of components of the complex resulted in the opposite effect (Wang et al., 2018). By expressing a mutant form of the E2 conjugase Ubc9, which results in strongly reduced activity of the sumoylation pathway, it was subsequently shown that under optimal growth conditions the net overall effect of sumoylation is to promote activity of the complex (Chymkowitch et al., 2017). This example shows that different sumoylation sites can have opposing effects on the activity of a protein complex, and that the cell targets specific sumoylation depending on the cellular state. We speculate that the same may be the case for the Ppar*γ* complex. Depending on environmental cues, cells use different sumoylation sites and SUMO isoforms to finetune the activity of the Ppar*γ* complex, resulting either in its activation or inhibition; however, we believe that during normal AD the overall effect of sumoylation on Ppar*γ* is to activate the complex. Clearly, more experiments are required to dissect the exact mechanism by which sumoylation regulates the Pparg complex.

In conclusion, our study provides novel insight into the role of the sumoylation pathway during dynamic changes in transcriptional regulation that occur during adipocyte differentiation, and also provides a wealth of data that should be of interest for researchers working in the fields of sumoylation, transcription and adipocyte differentiation.

## Acknowledgements

This work was supported by EPIC-XS (grant agreement EPIC-XS-823839, project number 0000062) funded by the Horizon 2020 programme of the European Union, a Helse Sør-Øst researcher grant (project number 2017064) and a Norwegian Research Council researcher grant (project number 301268) to PC. The work carried out in the Nielsen lab was in part supported by the Novo Nordisk Foundation Center for Protein Research, the Novo Nordisk Foundation (grant agreement numbers NNF14CC0001 and NNF13OC0006477), Danish Council of Independent Research (grant agreement numbers 4002-00051, 4183-00322A, 8020-00220B and 0135-00096B), and The Danish Cancer Society (grant agreement R146-A9159-16-S2). Sequencing was performed by the GenomEast platform, a member of the ‘France Génomique’ consortium (ANR-10-INBS-0009). This work was also supported in part by grants from the Norwegian Cancer Society (project number 182524), the Norwegian Research Council (261936) and the Research Council of Norway through its Centers of Excellence funding scheme, project number 262652. We thank Dr. Deo Prakash Panday for fruitful discussions during this project.

## Author contributions

Conceptualization, P.C.; Methodology, X.Z., I.A.H., A.N.P and P.C.; Formal Analysis, X.Z., I.A.H., S.L.G., T.Y., A.N.P and P.C.; Investigation, X.Z., I.A.H., A.N.P, G.F.L., L.R-A. and P.C.; Resources, A.K. and P.C.; Data Curation, X.Z., I.A.H., S.L.G. and A.N.P; Writing – Original Draft, P.C.; Writing – Review & Editing, X.Z., I.A.H., S.L.G., T.Y., B.J., J.M.E. M.L.N. and P.C.; Visualization, X.Z., I.A.H., S.L.G., T.Y., A.N.P and P.C.; Supervision, P.C.; Project Administration, P.C.; Funding Acquisition, P.C.

## STAR methods Lead Contact

Further information and requests for resources and reagents should be directed to and will be fulfilled by the corresponding author Pierre Chymkowitch.

## Materials Availability

This study did not generate new unique reagents.

## Data and code availability

The SLAMseq / Quantseq and SUMO ChIPseq datasets data have been deposited to the Gene Expression Omnibus with accession number GSE167222.

The accession numbers for published PPAR*γ* and RXR ChIPseq data are GSM340795, GSM340796, GSM340799, GSM340801, GSM340802 and GSM340805 (Nielsen et al., 2008).

The mass spectrometry proteomics data have been deposited to the ProteomeXchange Consortium via the PRIDE partner repository with the dataset identifier PXD024144.

## EXPERIMENTAL MODEL AND SUBJECT DETAILS

### 3T3-L1 culture, differentiation and treatments

3T3-L1 preadipocytes (CL-173, ATCC) were maintained in Dulbecco’s modified Eagle’s medium (DMEM) supplemented with 10% bovin calf serum (12138C, Sigma) and 1 % Penicillin/Streptavidin (P/S) at 37 °C in a humidified incubator in a 5 % CO_2_ in air atmosphere. Before adipogenic induction 3T3-L1 preadipocytes were grown to confluence by replacing the maintenance medium every second day for at least 4 days. Differentiation of 3T3-L1 cells was induced with differentiation medium (DMEM supplemented with 10 % fetal bovine serum (F7524, Sigma), 1mM dexamethasone (D4902, Sigma), 0.5 mM 3-isobutyl-1-methylxanthine (I5879, Sigma), 10 µg/mL insulin and 1 % P/S). At day 3 post adipogenic induction, the differentiation medium was replaced with the adipocyte maintenance medium (DMEM supplemented with 10 % FBS, 10 µg/mL of insulin and 1 % P/S). Adipocyte maintenance medium was then refreshed every second day. SUMOylation was inhibited by supplementing the culture medium with 0.5 µM of ML-792 (407886, Medkoo Biosite).

## METHOD DETAILS

### Lipid droplet staining

3T3-L1 cells were grown as described in “Adipocyte culture, differentiation and treatment”. At day 7 post induction cells were washed with PBS and fixed with 4.5 % formaldehyde during 10 minutes and rinsed 3 times with PBS. Lipid droplets were stained with a Bodipy 493/503 (D3922, Thermo Fisher Scientific) staining solution (1 µg/mL Bodipy 493/503, 150 mM NaCl) for 10 min at room temperature. Nuclei were stained with DAPI (0.5 µg/mL).

### Image acquisition

Images were acquired using the ImageXpress Micro Confocal device (Molecular devices, serial number 5150066). Widefield images were acquired using the 40 X S Plan Fluor ELWD objective as Z-series (step sizes 0.2-1 µm, depending on the staining). At least 4 representative fields per well were captured.

### Image analysis and quantification

All images were analyzed using the FIJI software (Fiji is just ImageJ) (National Institutes of Health, Bethesda, MD, USA). Z series images were corrected from bleaching then projected to obtain a “focused” single image corresponding to the different focal planes, using the *Stack Focuser* plugin (https://imagiej.nih.gov/ij/plugins/stack-focuser.html). Lipid droplets were segmented and quantified in single cell using the *Lipid Droplets Tool* macro (http://dev.mri.cnrs.fr/projects/imagej-macros/wiki/Lipid_Droplets_Tool) (Colitti et al., 2018).

### QuantSeq and SLAM-Sequencing sample preparation

3T3-L1 cells were grown as described in “*Adipocyte culture, differentiation and treatment*” and treated with DMSO or ML-792. Three biological replicates were harvested per condition. Prior harvesting, nascent RNAs were labelled with 100 µM of 4-thiouridine during 45 min. After labelling cells were lysed, homogenized and collected using 1 ml of RNAzol®RT (RN190-100, Molecular Research Center) before adding 0.4 ml of water per 1 ml of homogenate. The mixture was vortexed for 5 min and centrifuged at 12,000 g for 15 min. Following centrifugation, DNA, proteins and polysaccharides form a semisolid pellet at the bottom of the tube. The RNA remains soluble in the supernatant. The supernatant (1 ml maximum) was transferred to a new tube and 1 volume of 70% ethanol was added. The mixture was homogenized by pipetting (do not centrifuge). Samples were then applied to a RNeasy spin column (74104, Qiagen) and RNAs were purified according to Qiagen’s instructions. Base conversion was performed using the SLAMseq catabolic kinetics module (Lexogen, 062.24). RNA quantification and quality control were performed using Tape Station 4150 (Agilent).

### Library preparation and sequencing of QuantSeq and SLAMseq samples

mRNA-Seq libraries were generated according to manufacturer’s instructions from 500 ng of total RNA using the QuantSeq 3′mRNA-Seq Library Prep Kit for Illumina (FWD) (# 015, Lexogen GmbH, Vienna, Austria). Reverse transcription was initiated by oligo dT priming. After first strand cDNA synthesis the RNA was removed and second strand synthesis was initiated by random priming. Oligo dT primer and random primers contain Illumina-compatible adapter sequences. The resulting double-stranded cDNA was then purified and PCR amplified (30 sec at 98°C; [10 sec at 98°C, 20 sec at 65°C, 30 sec at 72°C] x 11 cycles; 1 min at 72°C), introducing i7 indexes. Surplus PCR primers were further removed by purification using SPRI-select beads (Beckman-Coulter, Villepinte, France) and the final libraries were checked for quality and quantified using capillary electrophoresis. The libraries were sequenced on Illumina Hiseq 4000 sequencer as Single-Read 50 base reads following Illumina’s instructions. Image analysis and base calling were performed using RTA 2.7.7 and bcl2fastq. 2.17.1.14. Adapter dimer reads were removed using DimerRemover (https://sourceforge.net/projects/dimerremover/).

### SLAMseq and QuantSeq data analysis

mm10 mouse genome assembly and Refseq 3’UTR coordinates were downloaded from UCSC (4 August 2020) using Table Browser. Sequencing reads were mapped and filtered with SlamDunk pipeline v0.4.3 (http://t-neumann.github.io/slamdunk/docs.html #document-Dunks). SlamDunk all (http://t-neumann.github.io/slamdunk/docs.html#all) was applied to full analysis for all samples. Reads with ≥ 1 T>C conversions were considered as labeled reads. Default settings for other parameters was followed.

Two parallel differential gene expression analyses were performed using DESeq2 R package (1.26.0). The total RNA reads were used for Quantseq analysis. The normalization size factor in Quantseq analysis was applied to global normalization for labeled reads. Time course experiments design (http://master.bioconductor.org/packages/release/workflows/vignettes/rnaseqGene/inst/d oc/rnaseqGene.html#time-course-experiments) was adapted for our data. Principal component analysis was performed after variance stabilizing transformation on total genes for both analyses.

### Chromatin immunoprecipitation (ChIP)

3T3-L1 cells were grown as described in “*Adipocyte culture, differentiation and treatment*” in 15 cm dishes. Two dishes were used per biological replicate and two biological replicates were collected for each time point.

Our ChIP procedure was adapted from (Baik et al., 2018). Cells were cross-linked in dish with 1% formaldehyde for 8 minutes. Formaldehyde was then neutralized using 125 mM glycine for 10 minutes. After two washes with cold PBS the cells were collected in the Lysis buffer (5 mM PIPES pH 7.5, 85 mM KCl, 0.5% NP40, 20 mM N-ethyl maleimide [NEM] and protease inhibitor cocktail [04693159001, Roche]) and incubated at 4°C for 10 minutes with rotation. Nuclei were centrifuged (1,500 rpm for 10 minutes at 4°C) and resuspended in a nucleus lysis buffer (50 mM Tris-HCl pH 7.5, 1% SDS, 10 mM EDTA, 20 mM NEM and protease inhibitor cocktail) and incubated at 4°C for 2 hours. Lysates were sonicated for 15 cycles (30 sec on / 30 sec off) at 4°C using a Bioruptor Pico sonicator (Diagenode). After sonication, lysates were centrifuged at 14,000 rpm for 10 minutes at 4°C. Protein concentration was assessed using the Bradford assay and 250 μg of chromatin were used for each immunoprecipitation. Input samples (12.5 μg) were saved. Samples were diluted 10-fold in the immunoprecipitation buffer (1.1% Triton X100, 50 mM Tris-HCl pH 7.5, 167 mM NaCl, 5 mM N-ethyl maleimide, 1 mM EDTA, 0.01% SDS, and protease inhibitor cocktail). Immunoprecipitations were carried out with 10 μg of SUMO-2/3 antibody (ab3742, Abcam) and 420 μl of Dynabeads Protein A (10001D, Thermo Fisher Scientific) overnight at 4°C. Beads were then washed 2 times in low-salt buffer (50 mM Tris-HCl pH 7.5, 150 mM NaCl, 1% Triton X100, 0.1% SDS, 1 mM EDTA), 2 times in high-salt buffer (50 mM Tris-HCl pH 7.5, 500 mM NaCl, 1% Triton X100, 0.1% SDS, 1 mM EDTA), 2 times in LiCl buffer (20 mM Tris-HCl pH 7.5, 250 mM LiCl, 1% NP40, 1% deoxycholic acid, 1 mM EDTA) and in TE buffer (10 mM Tris-HCl pH 7.5, 0.2% Tween20, 1 mM EDTA). Elution was done two time in 50 μL of 100 mM NaHCO3, 1% SDS at 65°C for 10 min under agitation. Chromatin cross-linking was reversed at 65°C for 5 hours with 280 mM NaCl and 88 μg/mL RNase DNase free (11119915001, Roche). Proteins were then digested using 88 μg/mL of Proteinase K (03115828001, Roche) during 1 hour at 65°C. DNA from immunoprecipitations and inputs were purified using the Qiagen PCR purification kit. DNA concentration was assessed using a Qubit device (Q32866, Invitrogen).

### Library preparation and sequencing of ChIP samples (ChIPseq)

ChIP samples were purified using Agencourt AMPure XP beads (Beckman Coulter) and quantified with the Qubit (Invitrogen). ChIPseq libraries were prepared from 10 ng of double-stranded purified DNA using the MicroPlex Library Preparation kit v2 (C05010014, Diagenode s.a., Seraing, Belgium), according to manufacturer’s instructions. In the first step, the DNA was repaired and yielded molecules with blunt ends. In the next step, stem-loop adaptors with blocked 5 prime ends were ligated to the 5 prime end of the genomic DNA, leaving a nick at the 3 prime end. The adaptors cannot ligate to each other and do not have single-strand tails, avoiding non-specific background. In the final step, the 3 prime ends of the genomic DNA were extended to complete library synthesis and Illumina compatible indexes were added through a PCR amplification (7 cycles). Amplified libraries were purified and size-selected using Agencourt AMPure XP beads (Beckman Coulter) to remove unincorporated primers and other reagents. The libraries were sequenced on Illumina Hiseq 4000 sequencer as Single-Read 50 base reads following Illumina’s instructions. Image analysis and base calling were performed using RTA 2.7.7 and bcl2fastq 2.17.1.14. Adapter dimer reads were removed using DimerRemover.

### ChIPseq sequencing data analysis

Reads were mapped to the Mus musculus genome (assembly mm10) using Bowtie (Langmead, 2010) v1.0.0 using the following parameters -m 1 --strata --best. Reads mapped in genomic regions flagged as ENCODE blacklist were removed (Amemiya et al., 2019). SUMO peaks were called with the ENCODE ChIPseq pipeline v1.3.6. Briefly, the pipeline ran quality controls and called peaks with spp v1.15.5 (Kharchenko et al., 2008). Reproductible peaks were kept after the IDR analysis was run (optimal IDR sets of peaks were kept). Peaks were annotated relative to genomic features using Homer annotatePeaks.pl v4.11.1 (Heinz et al., 2010). Known or *de novo* TF motifs were identified using HOMER findMotifsGenome.pl with default parameters. Heatmaps and mean profiles presenting read enrichments at various genomic locations were generated using Easeq software v1.111 (Lerdrup et al., 2016). To compare SUMO peaks enrichments over time, the union of all peak positions was computed with BEDtools v2.26.0 (Quinlan and Hall, 2010). Then, read counts per peak (union peak set) were normalized across libraries with the method proposed by Anders and Huber (Anders and Huber, 2010) and implemented in the Bioconductor package v1.24.0 (Love et al., 2014). Regions varying due to time effect were identified using a likelihood ratio test (LRT) with DESeq2. Resulting p-values were adjusted for multiple testing using the Benjamin and Hochberg method (Benjamini and Hochberg, 1995). Significant regions were those which adjusted p-value ≤ 0.05, absolute fold change > 1.5.

### RXR and Pparγ ChIPseq datasets

RXR and Ppar*γ* data were mapped to the Mus musculus genome (assembly mm10) using Bowtie (Langmead, 2010) v1.0.0 using the following parameters -m 1 --strata –best. Peaks were called using MACS2 callpeak with default parameters except for -g mm --nomodel --extsize 200. Peaks were annotated relative to genomic features using Homer annotatePeaks.pl v4.11.1 (Heinz et al., 2010).

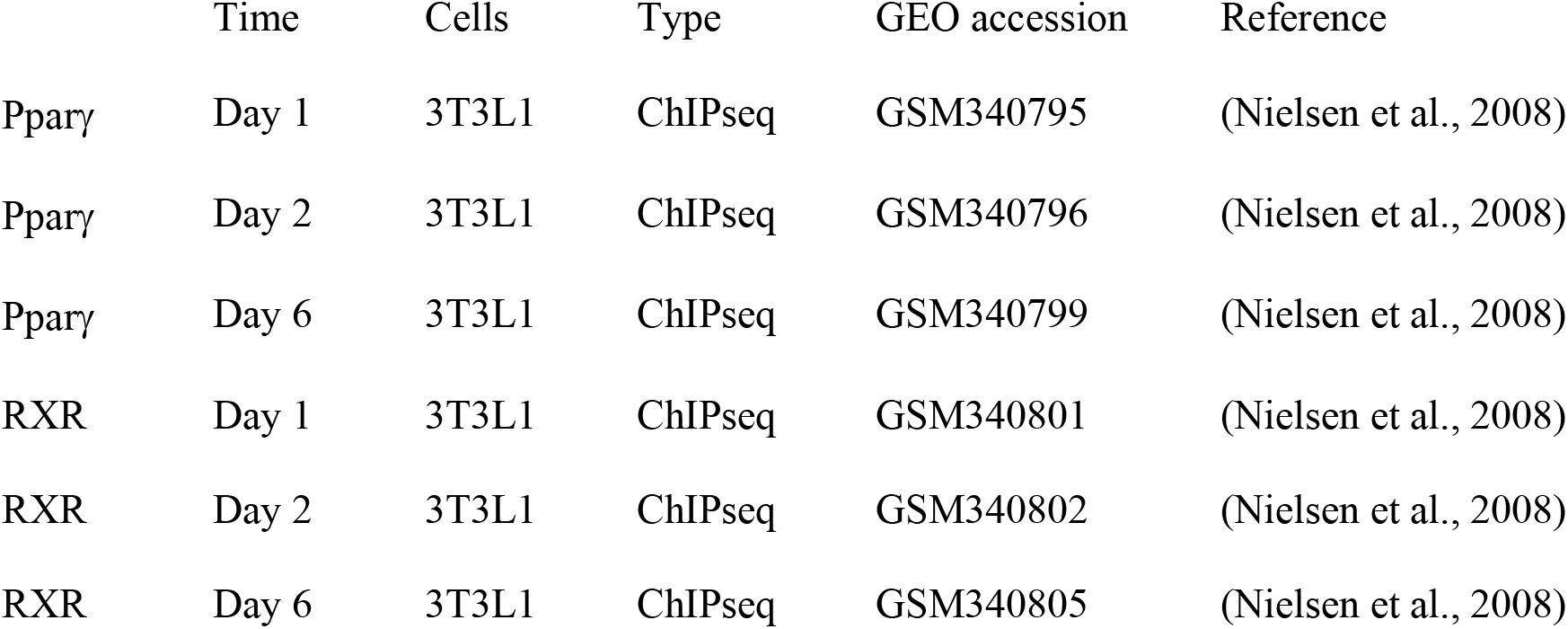

### SUMO Mass Spectrometry

3T3-L1 cells were grown as described in “*Adipocyte culture, differentiation and treatment*”. Four biological replicates were collected for each condition. Cells were washed with PBS supplemented with 20 mM NEM (E3876, Merck Life Science). Cells were vigorously lysed in guanidium lysis buffer (6M guanidine-HCl, 50 mM Tris pH 8.5, 20 mM NEM), after which they were immediately snap frozen. Lysates were stored at −80°C until further processing. In essence, sample preparation and SUMO-IP for native and endogenous mass spectrometry (MS) analysis was performed as described previously (Hendriks et al., 2018). Lysates were thawed at room temperature, after which they were supplemented with 5 mM chloroacetamide (CAA) and 5 mM Tris(2-carboxyethyl)phosphine (TCEP). Samples were homogenized via sonication using a microtip sonicator, at 30 W using three 10 s pulses, and afterwards cleared by centrifugation at 4,250*g*. Endoproteinase Lys-C (Wako) was added to samples in a 1:200 enzyme-to-protein ratio (w/w). Digestion was performed overnight, still, and at room temperature. Digested samples were diluted with three volumes of 50 mM ammonium bicarbonate (ABC), and a second round of overnight digestion was performed by addition of Lys-C in a 1:200 enzyme-to-protein ratio. Digests were acidified by addition of 0.5% trifluoroacetic acid (TFA), 1:100 vol/vol from a 50% TFA stock, after which they were transferred to 50 mL tubes and centrifuged at 4,250*g* and at 4°C for 30 min. Clarified digests were carefully decanted into clean 50 mL tubes, after which peptides were purified using C8 Sep-Pak cartridges (Waters) according to the manufacturer’s instructions. Sep-Pak cartridges with 500 mg C8 sorbent were used, with one cartridge used for each ∼25 mg of digested protein. Small and hydrophilic peptides were pre-eluted using 5 mL of 20% acetonitrile (ACN) in 0.1% TFA, and 3 mL of 25% ACN in 0.1% TFA. SUMOylated peptides were eluted using 1 mL of 35% ACN in 0.1 TFA, 1 mL of 40% ACN in 0.1% TFA, and 2 mL of 45% ACN in 0.1% TFA. For each replicate sample, all SepPak elutions were pooled in 50 mL tubes with small holes punctured into the caps, and then frozen overnight at −80°C. Deep-frozen samples were lyophilized to dryness for 48 h, with the pressure target set at 0.004 mbar and the condenser coil at −85°C.

### Crosslinking of SUMO antibody to beads

Overall, 750 μL of Protein G Agarose beads (Roche) were used to capture 400 μL of SUMO-2/3 antibody (8A2, acquired from Abcam, ab81371; ∼5-10 μg/μL antibody). All washing and handling steps were followed by centrifugation of the beads at 500*g* for 3 min in a swing-out centrifuge with delayed deceleration and careful aspiration of buffers, to minimize loss of beads. Beads were pre-washed 4 times with ice-cold PBS, split across three 1.5 mL tubes, after which the antibody was added and the tubes filled completely with ice-cold PBS. Beads and antibody were incubated at 4°C on a rotating mixer for 1 h, and subsequently washed 3 times with ice-cold PBS. Crosslinking of the antibody to the beads was achieved by addition of 1 mL of 0.2 M sodium borate, pH 9.0, which was freshly supplemented with 20 mM dimethyl pimelimidate (DMP). Crosslinking was performed for 30 min at room temperature on a rotating mixer, after which the crosslinking step was repeated once. SUMO-IP beads where then washed twice with ice-cold PBS, twice with 0.1 M glycine pH 2.8, and three times with ice-cold PBS, after which all beads were pooled in a single 1.5 mL tube and stored until use at 4°C in PBS supplemented with 10 mM sodium azide.

### Purification of SUMOylated peptides

Lyophilized peptides were dissolved in 10 mL ice-cold SUMO-IP buffer (50 mM MOPS, 10 mM Na2HPO4, 50 mM NaCl, buffered at pH 7.2) per 50 mg protein originally in the samples. Samples were clarified by centrifugation at 4,250*g* for 30 min at 4°C in a swing-out centrifuge with delayed deceleration. Samples were transferred to new tubes, after which 25 μL SUMO-IP beads was added per 50 mg protein originally in the samples. Samples were incubated at 4°C for 3 h in a rotating mixer, after which the beads were washed twice with ice-cold SUMO-IP buffer, twice with ice-cold PBS, and twice with ice-cold MQ water. Upon each first wash with a new buffer, beads were transferred to a clean 1.5 mL LoBind tube (Eppendorf). To minimize loss of beads, all centrifugation steps were performed at 500*g* for 3 min at 4°C in a swing-out centrifuge with delayed deceleration. Elution of SUMO peptides from the beads was performed by addition of 2 bead volumes of ice-cold 0.15% TFA, and performed for 30 min while standing still on ice, with gentle mixing every 10 min. The elution of the beads was repeated once, and both elutions were cleared through 0.45 μm spin filters (Millipore) by centrifuging at 12,000*g* for 1 min at 4°C. The two elutions from the same samples were pooled after clarification. Next, samples were pH-neutralized by addition of 1/10^th^ volume of 1 M Na_2_HPO_4_, and allowed to warm up to room temperature. Second-stage digestion of SUMOylated peptides was performed with 1 μg of Endoproteinase Asp-N (Roche). Digestion was performed overnight, at 30°C and shaking at 300 rpm, after which samples were frozen at −80°C until further processing.

### StageTip purification and high-pH fractionation of SUMO-IP samples

Preparation of StageTips (Rappsilber et al., 2003), and high-pH fractionation of SUMO-IP samples on StageTip, was performed essentially as described previously (Hendriks et al., 2018). Quad-layer StageTips were prepared using four punch-outs of C18 material (Sigma-Aldrich, Empore™ SPE Disks, C18, 47 mm). StageTips were equilibrated using 100 μL of methanol, 100 μL of 80% ACN in 200 mM ammonium, and two times 75 μL 50 mM ammonium. Samples were thawed out, and supplemented with 1/10^th^ volume of 200 mM ammonium, just prior to loading them on StageTip. The StageTips were subsequently washed twice with 150 μL 50 mM ammonium, and afterwards eluted as six fractions (F1-6) using 40 μL of 4, 7, 10, 13, 17, and 25% ACN in 50 mM ammonium. All fractions were dried to completion in LoBind tubes, using a SpeedVac for 2 h at 60°C, after which the dried peptides were dissolved using 10.5 μL of 0.1% formic acid.

### MS analysis

All samples were analyzed on EASY-nLC 1200 system (Thermo) coupled to a Q Exactive™ HF-X Hybrid Quadrupole-Orbitrap™ mass spectrometer (Thermo). For each run, 5 μL of sample was injected. Separation of peptides was performed using 15-cm columns (75 μm internal diameter) packed in-house with ReproSil-Pur 120 C18-AQ 1.9 µm beads (Dr. Maisch). Elution of peptides from the column was achieved using a gradient ranging from buffer A (0.1% formic acid) to buffer B (80% acetonitrile in 0.1% formic acid), at a flow of 250 nl/min. Gradient length was 80 min per sample, including ramp-up and wash-out, and an analytical gradient of 50 min. The buffer B ramp for the analytical gradient was as follows: F1: 13-24%, F2: 14-27%, F3-5: 15-30%, F6: 17-32%. The columns were heated to 40°C using a column oven, and ionization was achieved using a Nanospray Flex Ion Source (Thermo) with the spray voltage set at 2 kV, an ion transfer tube temperature of 275°C, and an RF funnel level of 40%. Full scan range was set to 400-1,600 *m*/*z*, MS1 resolution to 60,000, MS1 AGC target to 3,000,000, and MS1 maximum injection time to 60 ms. Precursors with charges 2-6 were selected for fragmentation using an isolation width of 1.3 *m*/*z*, and fragmented using higher-energy collision disassociation (HCD) with normalized collision energy of 25. Precursors were excluded from re-sequencing by setting a dynamic exclusion of 60 s. MS2 resolution was set to 60,000, MS2 AGC target to 200,000, minimum MS2 GC target to 20,000, MS2 maximum injection time to 120 ms, and loop count to 7.

### Analysis of MS data

All MS RAW data was analyzed using the freely available MaxQuant software, version 1.5.3.30 (Cox and Mann, 2008). All data was processed in a single computational run, and default MaxQuant settings were used, with exceptions specified below. For generation of the theoretical spectral library, the mouse FASTA database was downloaded from Uniprot on the 14^th^ of February, 2020. The mature sequence of SUMO2 was inserted in the database to allow for detection of free SUMO. In silico digestion of theoretical peptides was performed with Lys-C, Asp-N, and Glu-N, allowing up to 8 missed cleavages. Variable modifications used were protein N-terminal acetylation, methionine oxidation, peptide N-terminal pyroglutamate, Ser/Thr/Tyr phosphorylation (STY), and Lys SUMOylation, with a maximum of 3 modifications per peptide. The SUMO mass remnant was defined as described previously (Hendriks et al., 2018); DVFQQQTGG, H_60_C_41_N_12_O_15_, monoisotopic mass 960.4301, neutral loss b7-DVFQQQT, diagnostic mass remnants [b2-DV, b3-DVF, b4-DVFQ, b5-DVFQQ, b6-DVFQQQ, b7-DVFQQQT, b9-DVFQQQTGG, QQ, FQ, FQQ]. Label-free quantification was enabled, with “Fast LFQ” disabled. Maximum peptide mass was set to 6,000 Da. Stringent MaxQuant 1% FDR filtering was applied (default), and additional automatic filtering was ensured by setting the minimum delta score for modified peptides to 20, with a site decoy fraction of 2%. Second peptide search was enabled (default). Matching between runs was enabled, with a match time window of 1 min and an alignment window of 20 min. For protein quantification, the same variable modifications were included as for the peptide search. To further minimize false-positive discovery, additional manual filtering was performed at the peptide level. All modified peptides were required to have a localization probability of >75%, be supported by diagnostic mass remnants, be absent in the decoy database, and have a delta score of >40 in case SUMO modification was detected on a peptide C-terminal lysine not preceding an aspartic acid or glutamic acid. All multiply-SUMOylated peptides were discarded, unless the corresponding SUMO sites were also identified by singly-SUMOylated peptides. SUMO target proteins were derived from the “proteinGroups.txt” file, and all post-filtering SUMO sites were manually mapped. Only proteins containing at least one SUMO site were considered as SUMO target proteins, and other putative SUMO target proteins were discarded.

### Calculation of SUMO density and equilibrium

SUMO density was calculated by dividing the total sum of all SUMO site intensity (in arb. units., corresponding to ion current) by the amount of total protein starting material (in mg). This calculation was performed for each replicate separately, and visualized as an average ± standard deviation. The mature sequence of SUMO2 was included as a FASTA file in the MaxQuant search, to allow detection of free mature SUMO2/3. For quantification of the SUMO equilibrium, the “modificationSpecificPeptides.txt” MaxQuant output file was used, and all peptides modified by SUMO2/3, and peptides derived from SUMO2/3 itself, were considered. Modification of any SUMO family member by SUMO 2/3 was considered chain formation, with all other conjugation considered as global target modification. Peptides derived from SUMO2/3 were sub-classed as internal, mature free SUMO2/3, immature SUMO2, or immature SUMO3. Peptides ending in QQTGG (predominantly DVFQQQTGG) were considered as mature free SUMO2/3. Peptides containing but not ending with QQTGG were considered as immature SUMO2 (DVFQQQTGGVY), or immature SUMO3 (DVFQQQTGGSASRGSVPTPNRCP).

### Western blotting

3T3-L1 cells were grown as described in “*Adipocyte culture, differentiation and treatment*”. Cells were washed with PBS prior lysis in ice-cold RIPA buffer (150 mM NaCl, 50 mM Tris-HCl pH 7.5, 5 mM EDTA, 1 % NP-40, 0.5 % Na-deoxycholate, 0.1 % SDS) freshly supplemented with protease inhibitor (Roche) and 20 mM of NEM. Lysates were sonicated 5 min (30 sec on, 30 sec off) then centrifuged 15 min at 14 000 rpm and 4 °C. To eliminate lipids supernatants were applied to a RNeasy column and centrifuged at 10,000 rpm and 4 °C for 1 min (Qiagen, 74104). The flow through was collected and protein concentrations were assessed using the Bradford assay. 30 µg of proteins were used for each western blot and proteins were detected using anti-SUMO2/3 (ab3742, Abcam) and anti-TBP (ab51841, Abcam) antibodies.

### Data analysis

The hierarchical clustering and heatmaps were generated using Perseus software v1.6.10.50 (Tyanova et al., 2016). For SUMO2/3 proteomics, the label-free quantified (LFQ) value for four bio-replicates was averaged. The averaged value was log2 transformed, imputed with default setting, and normalized by row Z score transformation. For SLAMseq data, the normalized count for three bio-replicates was averaged. The averaged normalized count was log2 transformed and normalized by row Z score transformation. The row dendrogram was generated based on Euclidean distance and processed with k-means. Z score was used to generate cluster profiles in parallel.

GO analysis was performed using clusterProfiler R package (Yu et al., 2012). The statistical significance was specified as adjusted p value < 0.05. Redundant GO terms were simplified according to similarity measured with “Wang” method.

The scatter plot was generated using Perseus software. Log2 transformed normalized counts were plotted. Meanwhile, Pearson correlation coefficient and P value were calculated. Venn diagram was generated using VennDiagram R package (https://CRAN.R-project.org/package=VennDiagram).

For each single gene, Pearson correlation coefficient was calculated by comparing time-course profiles in different datasets in Excel. The density plot for the distribution of Pearson correlation coefficient was generated using EaSeq software v1.111 (Lerdrup et al., 2016). The ggplot2 R package v3.3.2 (https://ggplot2.tidyverse.org) was used to generate box, histogram, bar, dot and line plots in the study.

Integrative Genomics Viewer (Robinson et al., 2011) was used for SLAMseq and ChIPseq data visualization with normalized bigWig files.

### Quantification and statistical analysis

Statistical analysis was performed using GraphPad Prism (v7). For immunofluorescence data, Results are presented as mean ± SD. Between groups, statistical significance was calculated using unpaired, two tailed Student’s t tests. The significance of mean comparison in boxplots for time course studies was analyzed using stat_compare_means function in R. The paired t-test method was specified. Non-significant: p > 0.05, *: p ≤ 0.05, **: p ≤ 0.01, ***: p ≤ 0.001, ****: p ≤ 0.0001. P values < 0.05 were considered as significant. For overrepresentation analysis, Hypergeometric test was used for testing significance in R. P values were corrected by Benjamini-Hochberg method.

---

**Fig. S1.**
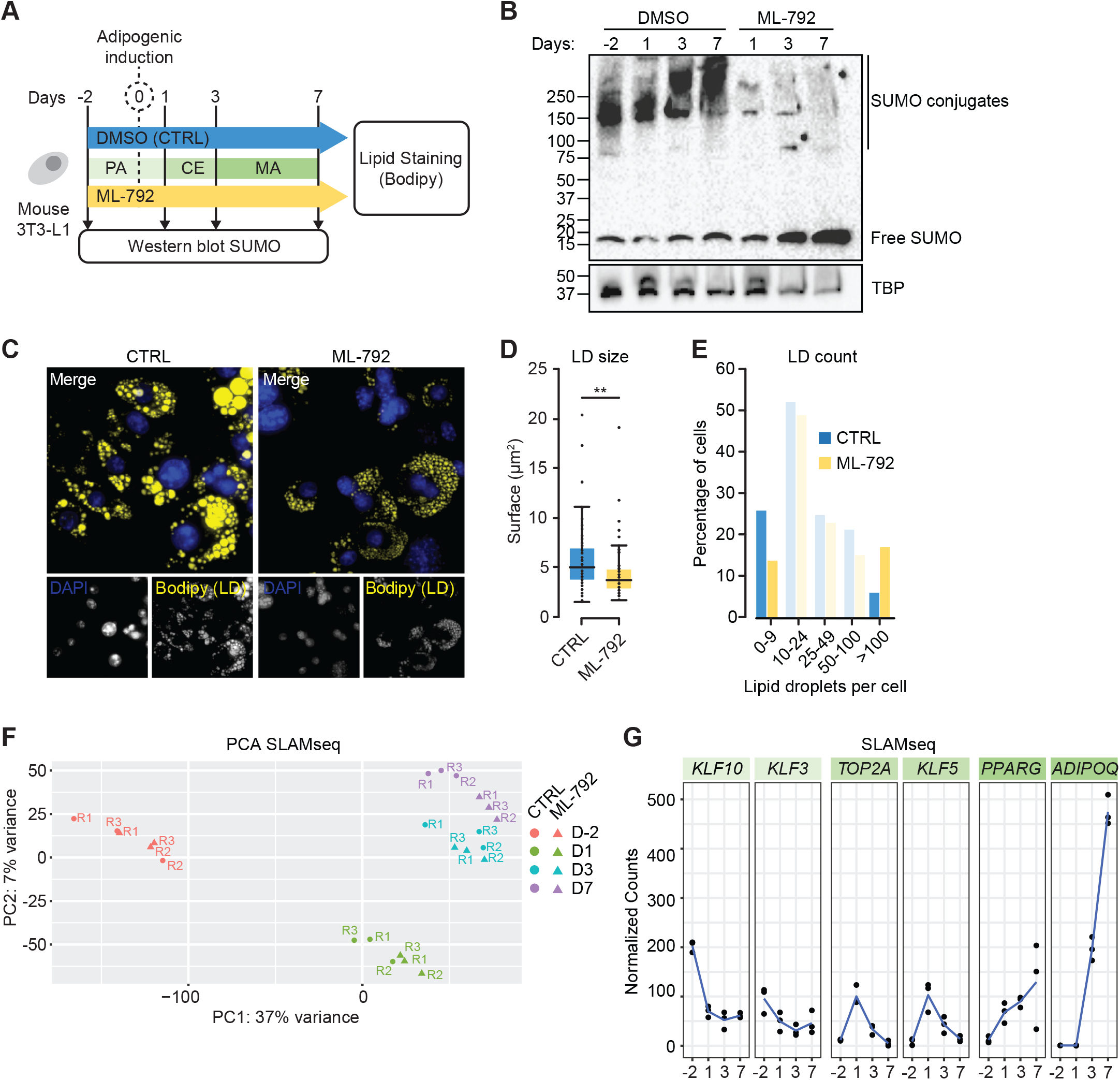
ML-792 triggers lipoatrophy. (A) Experimental layout of time-course experiments. (B) Western blot analysis of SUMO2/3 during adipogenesis in DMSO and ML-792-treated 3T3-L1 cells. Cells treated with DMSO or ML-792 (0.5μM) were collected at indicated time points. Cell lysates were subjected to western blot using indicated antibodies. TBP was used as loading control. (C) Immunofluorescence analysis of DMSO or ML-792 treated adipocytes (Day 7). Lipid droplets were stained with lipid probe Bodipy. Nucleus was counterstained with DAPI. Images were acquired using the ImageXpress Micro Confocal device (40x). Images are representative of at least 3 biological replicates. (D) The size of lipid droplets in DMSO or ML-792 treated adipocytes (Day 7) was measured using FIJI software. The surface area is measured in µm^2^ (y axis). At least 4 representative fields per well were analyzed. The significance of mean comparison is determined by unpaired t-test. ns: p > 0.05, *: p ≤ 0.05, **: p ≤ 0.01, ***: p ≤ 0.001, ****: p ≤ 0.0001. (E) The percentage of adipocytes (y axis) in subpopulations categorized based on lipid droplet numbers per cell (x axis). (F) PCA of SLAMseq data. (G) Profiles of representative transcripts in time-course modules in SLAMseq. The y axis is presented as normalized counts.

**Fig. S2.**
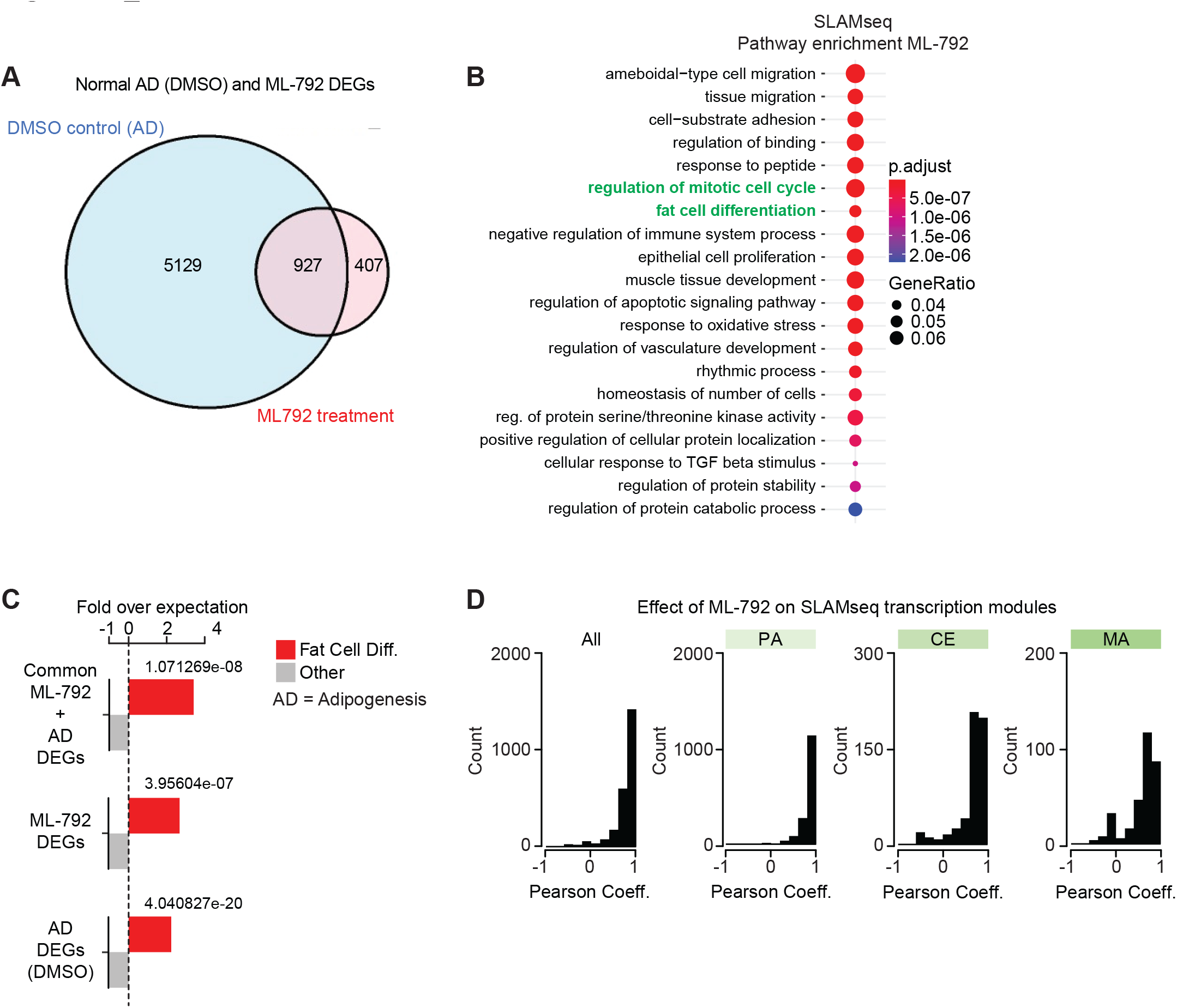
ML-792 affects adipogenesis transcription modules. (A) Venn diagram of control (DMSO) time-course DE transcripts and ML-792 treatment DE transcripts. (B) GO analysis of ML-792 treatment DEGs in SLAMseq. (C) Enrichment of fat cell differentiation term in DEG subsets defined in *(A)*. (D) Distributions of Pearson correlation coefficients in transcription modules. P: pattern effect, A: amplitude effect. Y axis in *(C)* is represented as transcript count numbers. The y axis, presented as Pearson correlation coefficient, is segmented into 10 bins. The number of transcripts within each bin is presented as a color code.

**Fig. S3.**
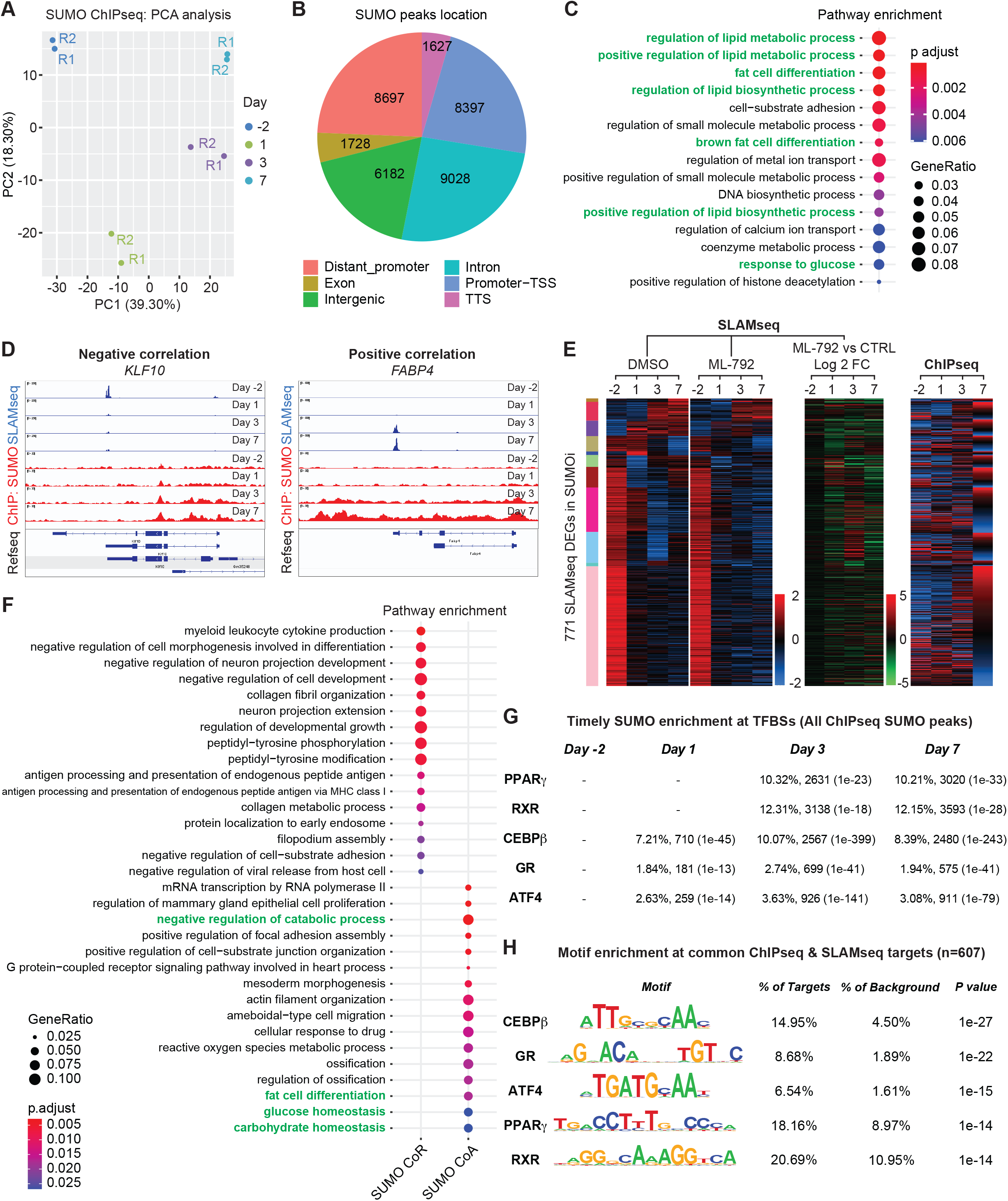
Characterisation of the SUMO chromatin landscape of differentiating 3T3-L1 cells. (A) PCA of SUMO ChIPseq data. (B) Genome-wide distribution of SUMO peaks. (C) GO analysis of target genes with differential binding SUMO peaks over the time-course. (D) Snapshot from the IGV genome browser for ChIPseq and SLAMseq data at *KLF10* (negative correlation) and *FABP4* (positive correlation) loci. (E) Heatmaps of transcriptions in DMSO (see Fig. 1D), ML-792 and SUMO ChIPseq during AD. Log2 fold change between ML-792 and DMSO treatments were calculated. (F) GO analysis of genes at which SUMO acts as corepressor (CoR) and coactivator (CoA). (G) Bimodal test of TFBS enrichment of SUMO binding across time points as observed in Fig. 3L. (H) Motif enrichment of SUMO binding at TFBSs of SLAMseq DEGs *(F* and *G)*.

**Fig. S4.**
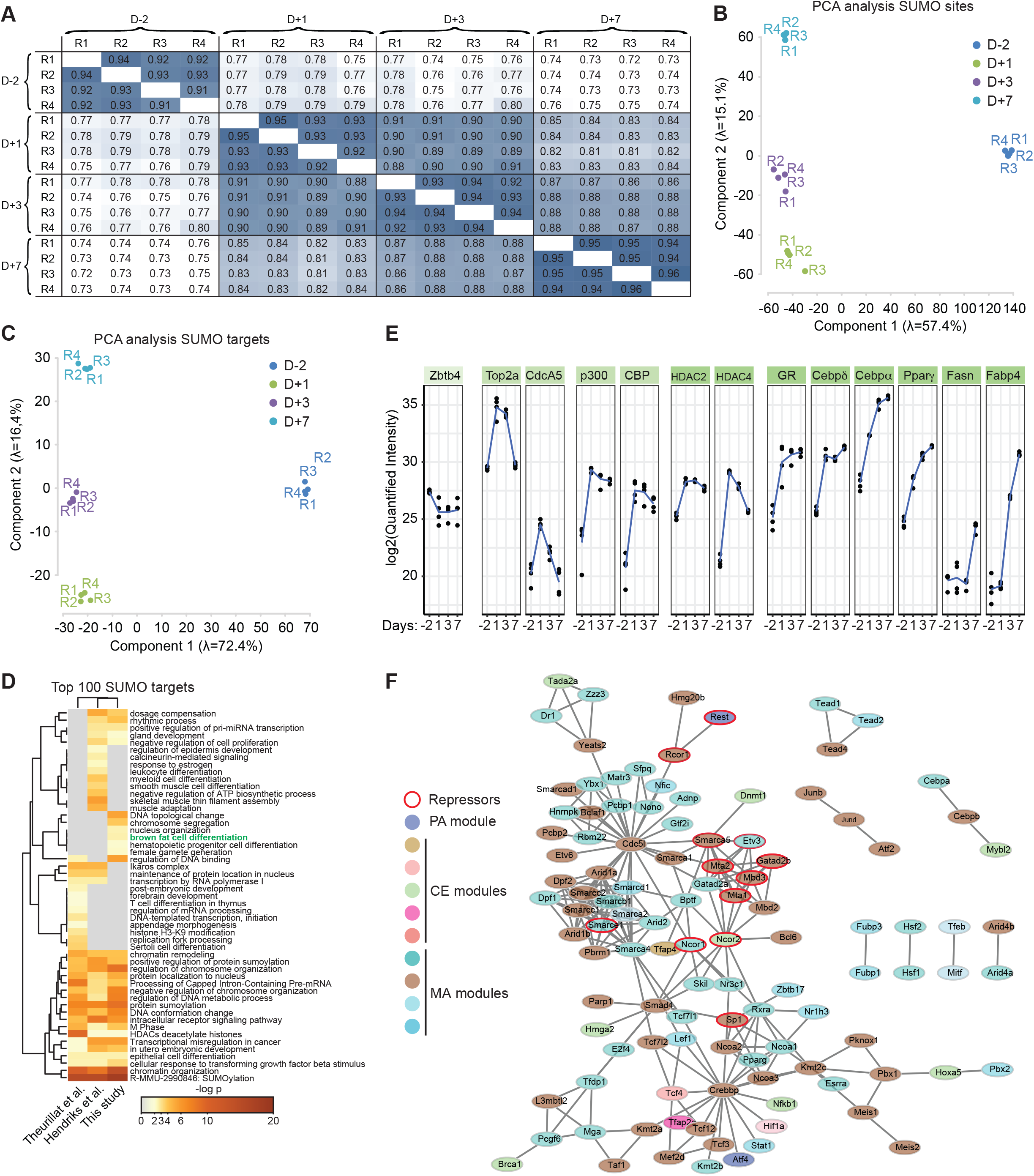
Site specific characterization of the endogenous SUMOylome of differentiating adipocytes. (A) Pearson correlation analysis of MS bioreplicates. (B) PCA of site-specific sumoylation across samples. (C) PCA of SUMO targets. (D) Metascape analysis of the top 100 SUMO targets from the studies. (E) Sumoylation profiles of representative SUMO targets in AD sumoylation modules. (F) STRING network analysis of sumoylated TFs. Node color: same as the colors of sumoylation module; Node border: red indicates transcriptional repressor. (G) STRING network of the Pparγ/RXR TF complex. (H) Loess regression lineplot of sumoylation intensity for the main factors in Pparγ/RXR complex. Solid line: TFs and co-activators; dashed line: co-repressors. Z scores are calculated for log2 transformed label-free quantified (LFQ) values.

**Fig. S5.**
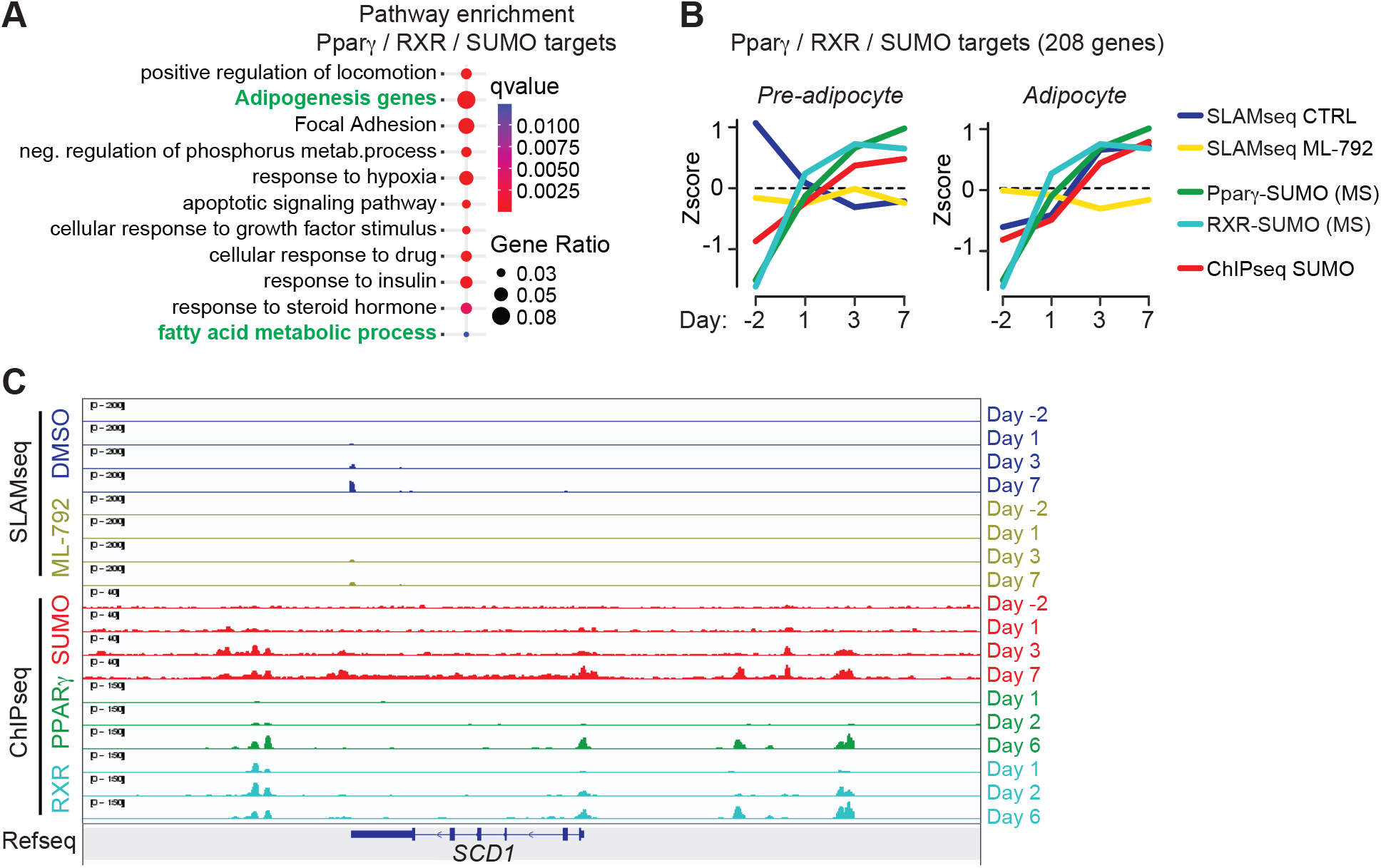
The Pparγ, RXR and SUMO regulon. (A) GO analysis of Pparγ, RXR and SUMO common targets. (B) Loess regression lineplots showing transcription profiles (CTRL, blue; ML-792, yellow), SUMO binding at PPRE (red) and SUMOylated Pparγ (green) and RXR (Turquoise) at Pparγ, RXR and SUMO common target genes. Z scores are calculated for log2 transformed normalized counts (transcript or SUMO binding site) or log2 transformed label-free quantified (LFQ) values (SUMO proteomics). (C) Snapshot of IGV genome browser showing the activation of the Pparγ/RXR target gene *SCD1* during AD (SLAMseq, DMSO), the downregulation of *SCD1* after ML-792 treatment (SLAMseq, ML-792) and the coordinated recruitment of SUMO, Pparγ and RXR at *SCD1* during AD (ChIPseq).

